# Large-scale CRISPRi and transcriptomics of *Staphylococcus epidermidis* identify genetic factors implicated in commensal-pathogen lifestyle versatility

**DOI:** 10.1101/2021.04.29.442003

**Authors:** Michelle Spoto, Johanna P. Riera Puma, Elizabeth Fleming, Changhui Guan, Yvette Ondouah Nzutchi, Dean Kim, Julia Oh

**Affiliations:** The Jackson Laboratory for Genomic Medicine, Farmington, Connecticut, USA; The University of Connecticut Health Center, Farmington, Connecticut, USA

**Keywords:** CRISPRCas9, large-scale knockdown, CRISPRi, transcriptomics

## Abstract

*Staphylococcus (S.) epidermidis* is a ubiquitous human commensal skin bacterium that is also one of the most prevalent nosocomial pathogens. The genetic factors underlying this remarkable lifestyle plasticity are incompletely understood, much due to the difficulties of genetic manipulation, precluding high-throughput functional profiling of this species. To probe *S. epidermdis’* versatility to survive across a diversity of skin sites and infection niches, we developed a large-scale CRISPR interference (CRISPRi) screen complemented by transcriptional profiling (RNA-seq) across 24 diverse environmental conditions and piloted a droplet-based CRISPRi approach to enhance throughput and sensitivity. We identified putative essential genes, importantly, revealing amino acid metabolism as crucial to survival across diverse environments and demonstrated the importance of trace metal uptake for survival under multiple stress conditions. We identified pathways significantly enriched and repressed across our range of stress and nutrient limited conditions, demonstrating the considerable plasticity of *S. epidermidis* in responding to environmental stressors. We postulate a mechanism by which nitrogen metabolism is linked to lifestyle versatility in response to hyperosmotic challenges, such as those encountered on human skin. Finally, we examined *S. epidermidis* survival under acid stress and hypothesize a role for cell wall modification as a vital component of the survival response in acidic conditions. Taken together, this study integrates large scale CRISPRi and transcriptomics data across multiple environments to provide insights into a keystone member of the human skin microbiome. Our results additionally provide a valuable benchmarking analysis for CRISPRi screens and are rich resource for other staphylococcal researchers.

**Author summary:** *Staphylococcus epidermidis* is an important bacteria of the skin microbiome. While it has an important role in skin health, it can also be a major infectious agent, especially in bloodstream and catheter infections. Understanding the underlying genes and pathways that contribute to *S. epidermidis’* ability to have both health and disease-associated abilities will be important to promoting the former and targeting the latter. Yet the function of many *S. epidermidis* genes, particularly in skin and infection environments, remains unknown. We developed a CRISPRi platform to knock down the function of *S. epidermidis* genes to better understand to what degree they are essential for growth in these environments. We complemented this gene essentiality data with gene expression data in the same environments to understand how regulation of these genes contribute to *S. epidermidis’* survival. These large-scale data generated numerous hypotheses for new genetic links to *S. epidermidis’* growth versatility.

## Introduction

Sophisticated and accessible sequencing technologies and computational tools to reconstruct complex metagenomes have greatly expanded the catalog of microbial species associated with a wide variety of different ecosystems. Understanding how these different species contribute to their respective ecosystems is one of microbial ecology’s next great challenges.

For this, experimental investigations at the gene level are vital to studying the behavior of any individual organism. Computational approaches leveraging homology to known sequences in model organisms are useful for predicting gene function, but species and strain-specific differences and erroneous or sparse gene annotations preclude a more extensive reliance on computational predictions. However, gold-standard approaches for characterizing gene function, such as gene knockouts, are laborious to create and phenotype on a scale that would allow a similarly rapid and comprehensive investigation of gene function in non-model organisms.

With the advent of large-scale methods for gene fitness investigation, such as transposon mutagenesis sequencing(van Opijnen et al., 2009) and antisense RNA inhibition (Forsyth et al., 2002), high throughput gene function screens have provided a wealth of fitness data for an increasing diversity of microbes in different growth conditions (Carey et al., 2018; Coe et al., 2019; Salama et al., 2004) . But even with these methods, these high-throughput investigations could still be impractical for non-model microbes, especially those intractable to genetic manipulation. For example, restriction-modification systems may prevent the introduction of foreign DNA, or homologous recombination efficiency may be too low to enable direct modification of chromosomal loci. Fortunately, rapid developments in CRISPR-Cas9 technology have provided a new, powerful toolkit for gene modification and the discovery of gene function (Jiang et al., 2013).

In particular, the development of CRISPR interference (CRISPRi)(Gilbert et al., 2014; Qi et al., 2013), has opened a new avenue for large-scale studies of gene function in prokaryotes that does not rely on direct genetic manipulation(Cui et al., 2018; Lee et al., 2019; Peters et al., 2016; Rousset et al., 2018a; Wang et al., 2018). CRISPRi is a modification of the CRISPR-Cas system that employs a catalytically inactive Cas9 protein (dCas9) for transcriptional repression. Rather than inducing a double stranded break at a target sequence – which can be lethal in prokaryotes that have limited repair mechanisms – the dCas9 protein, guided by the targeting guide RNA, inhibits transcription of the target sequence via steric hinderance (Bikard et al., 2013). In prokaryotes, for example, CRISPRi has been used for gene essentiality screening (Lee et al., 2019; Wang et al., 2018), to map non-coding RNAs to growth phenotypes (Wang et al., 2018), and to investigate metabolic functions(Donati et al., 2021).

Here, we apply CRISPRi to the high throughput gene fitness screening of a keystone skin commensal, non-model organism, *Staphylococcus (S.) epidermidis.* We and others have shown that *S. epidermidis* has key roles in maintaining the integrity and health of the skin microbiome, for example, through modulation of the immune system (Lai et al., 2009; Linehan et al., 2018; Naik et al., 2012; Scharschmidt et al., 2015), the prevention of colonization by more virulent pathogens(Cogen et al., 2010a, 2010b; Lai et al., 2010) such as *S. aureus,* and suppression of virulence (Zhou et al., 2020). However, as a common cause of bloodstream and medical device infections, *S. epidermidis* is also implicated in skin disease (Byrd et al., 2017; Williams et al., 2020) and is a leading nosocomial pathogen (National Nosocomial Infections Surveillance System, 2004) prolonging hospital stays and increasing patient morbidity (Gray et al., 1995; Molina et al., 2013). Yet it is commonly assumed that these *S. epidermidis* infections are seeded from patients’ own skin, where it is ubiquitous (Oh et al., 2014, 2016) and surprisingly diverse within an individual, with unique skin site specification and functional niche adaptation (Zhou et al., 2020). We speculate that its genetic diversity enables this lifestyle versatility as a ubiquitous skin colonizer and opportunistic pathogen.

Despite its importance in skin and infectious disease, far less is understood about *S. epidermidis’* growth and survival than its more virulent cousin, *S. aureus*. Indeed, nearly 40% of genes lack adequate annotation due to limited genetic manipulation of *S. epidermidis* to date. Indeed, due to an active restriction-modification system (Corvaglia et al., 2010; Waldron and Lindsay, 2006) and a limited ability to undergo homologous recombination, classical techniques like gene knockout or transposon mutagenesis are technically challenging and time-consuming to implement in *S. epidermidis* (Widhelm et al., 2014). As a human-health associated microbe and a difficult-to-study non-model organism, *S. epidermidis* is an excellent candidate for high-throughput gene fitness screening with CRISPRi.

Here, we generated the first high-throughput investigation of gene fitness in *S. epidermidis* using a large-scale CRISPRi knockdown pool of ∼14k strains. We screened this pool across 24 diverse stress and nutrient-limiting conditions selected to identify genes that could contribute to *S. epidermidis’* diverse metabolic needs for its lifestyle versatility. To further probe gene function, we complemented our fitness analysis with RNA-seq on a subset of 18 of these conditions to investigate the transcriptomic adaptation to these environments. Overall, we identified putative essential genes required for *S. epidermidis* survival across varied environmental landscapes, with amino acid metabolism, ion uptake, and nitrate/nitrite reduction genes particularly implicated in its lifestyle versatility. We also comment on potential limitations of CRISPRi in non-model organisms given inherent species-specific characteristics, and pilot droplet-based CRISPRi for increased throughput and sensitivity. This dataset represents a large-scale demonstration of CRISPRi in a non-model staphylococcal species, and is a resource to staphylococcal researchers in its scope characterizing gene fitness and expression across a diversity of environmental conditions.

## Results

### Pooled CRISPRi can identify known essential genes

We previously reported our CRISPRi/Cas targeting vector, which contains the necessary CRISPR/Cas machinery for CRISPRi, including dCas9 under the control of an anhydrotetracycline (aTc) inducible promoter and a combined tracrRNA/crRNA (Spoto et al., 2020). We validated this vector by generating individual knockdown strains of putative essential genes in *S. epidermidis* and identifying strong growth defects (Spoto et al., 2020). Here, we confirmed that this CRISPRi vector was suitable for identifying essential genes in a pooled screen. We generated small ‘mock’ pools with four strains (targeting 2 known essential and 2 nonessential genes), then 500 strains (both essential and non-essential). We used log2 fold change (log2FC, defined as the log2 of the fold change of the normalized read counts of a guide in the final pool / initial pool) as a proxy for guide knockdown efficiency to analyze the relationship of our guide features to knockdown; this approach has been previously used to study CRISPRi targeting in prokaryotes (Cui et al., 2018). We found that the majority of guide “hits” were to essential genes (volcano plot depicting false discovery rate (fdr)-adjusted p-values vs. log2FC, Fig. 1), thus supporting the feasibility of our approach.

**Figure 1.**
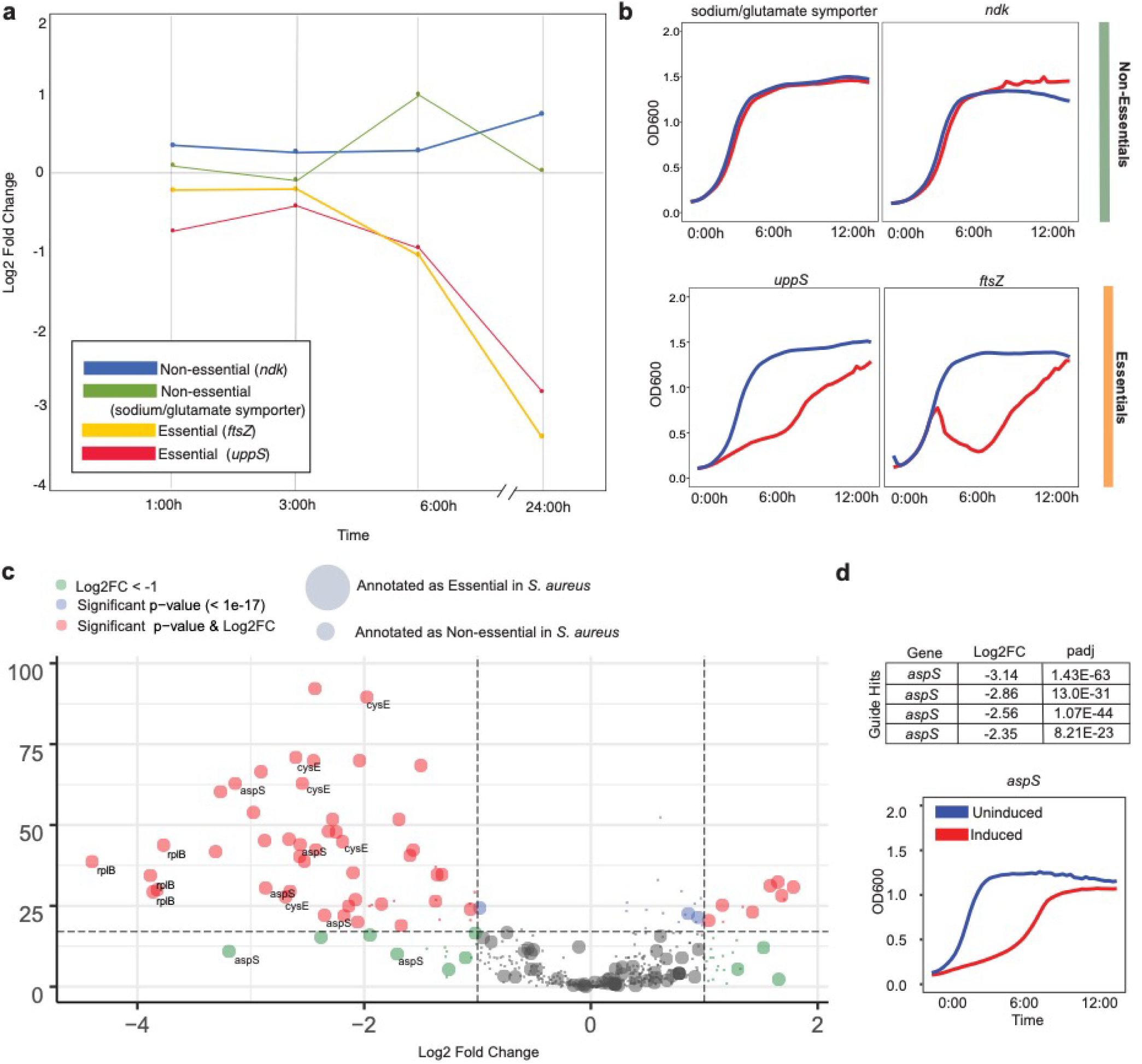
Pooled CRISPRi identifies known essential genes. A) Line chart depicting log2FC (final pool / initial pool) of essential and non-essential gene knockdowns across a 24 hour growth assay. B) Individual growth assays across 16 hours for each of these 4 strains depicting growth defects for essential gene knockdowns. C) Volcano plot of FDR-adjusted p-value (“padj”) and log2FC values for the pool of ∼ 500 knockdown strains. Guides with a significant padj value and a log2FC < -1 are depicted in red. Guides targeting known essential genes are depicted as large circles; known non-essential genes are depicted as small circles. All guides targeted to a select number of known essential genes (*rplB, cysE, aspS*) are shown, depicting that some genes are identified by multiple guides. D) log2FC and padj values for guide-hits to *aspS* are depicted alongside an individual growth curve for an *aspS* gene knockdown strain. Similar growth curves for *cysE* and *rplB* (and others) can be found in Spoto et al., 2020.

### Large-scale CRISPRi is effective for identifying gene fitness defects in rich media

#### Guide design and pool creation

Using a tiling approach to comprehensively target the *S. epidermidis* genome, ∼60k guides were designed by a customized version of our design script (Spoto et al., 2020) (STable 1). For our proof of principle, we selected *S. epidermidis* strain Tü3298 (Moran and Horsburgh, 2016), for its transformability (Winstel et al., 2016). Versus a gene-based approach for guide design, a tiling approach additionally covers protein coding sequences on the non-template strand, which has been shown to be an effective knockdown strategy for CRISPRi in prokaryotes, as well as other genomic elements, e.g., unknown promoter regions, rRNA and tRNA sequences, repeat regions, and others (Wang et al., 2018). Guides were cloned into a previously validated shuttle vector (Spoto et al., 2020). Following microarray-based guide synthesis, amplification, cloning in *E. coli* and finally, phagemid transformation into *S. epidermidis*, a loss of guides at each step resulted in a final knockdown library containing 14,341 unique guides targeting 1608 genes (∼67% of Tü3298 predicted protein coding sequences) (Fig. 1a). The number of guides per gene ranged from 1-31, depending on gene length and GC content (median 2 guides per gene). GC content of guides ranged from 35-85% (median 42%). The most pronounced limitations to guide design and final pool composition were the low GC content of the *S. epidermidis* genome (33%, and we set a threshold of at least 35% GC content for our guides, in line with previously published reports) and the multi-step transformation in *S. epidermidis* with a bottle neck at the intermediate *S. aureus* transformation stage (additional results on guide design is in Supplemental Notes). We note that improvements to transformation efficiency are a crucial step in the generalizability of large-scale CRISPR-Cas screens to non-model microbes.

#### CRISPRi screening in rich media is reproducible and can discriminate essential vs non-essential genes

We characterized the reproducibility of our CRISPRi knockdown library to identify strains with growth defects, using rich media (tryptic soy broth (TSB)) as our baseline. As anticipated, induction of dCas9 expression by aTc affected the overall composition of the knockdown library. Principle coordinate analysis (PCA) of normalized read counts (Fig. 1d) differentiated no-knockdown (uninduced) samples while knockdown (induced) samples clustered together by time point (Fig. 1d), indicating increasing knockdown phenotype over longer growth periods. The composition of the knockdown library was highly correlated between technical and biological replicates (Fig. 1b, Pearson correlation of normalized read counts, R=0.95 and R=0.98 respectively.) Despite high correlation between biological replicates, each screening assay was performed at least in triplicate to improve confidence in the final screen results. Additionally, the distribution of log2FCs for predicted essential (predicted from the literature) and predicted non-essential genes demonstrated a shift toward negative for guides targeting putative essential genes as compared to putative non-essential genes (Fig. 1c). Taken together, these results confirm the feasibility of using our large-scale knockdown pool for robust identification of genes that affect growth *in vitro*.

### CRISPRi screening identifies putative essential and stress response genes across multiple conditions

After initial characterizations in rich media, we sought to extend our gene phenotyping to different environments. 24 *in vitro* conditions (Table 1) were selected with the goal of providing insights into *S. epidermidis’* basic physiological response to stress and growth in different facets of skin and infection-related conditions. For example, genes required for *S. epidermidis* survival across the range of environmental stressors encountered during infection may present additional therapeutic targets not identifiable through screening of gene function in rich media. First, we sought to identify genes essential for growth irrespective of environmental condition and genes vital for growth in multiple stress conditions.

**Table 1.**
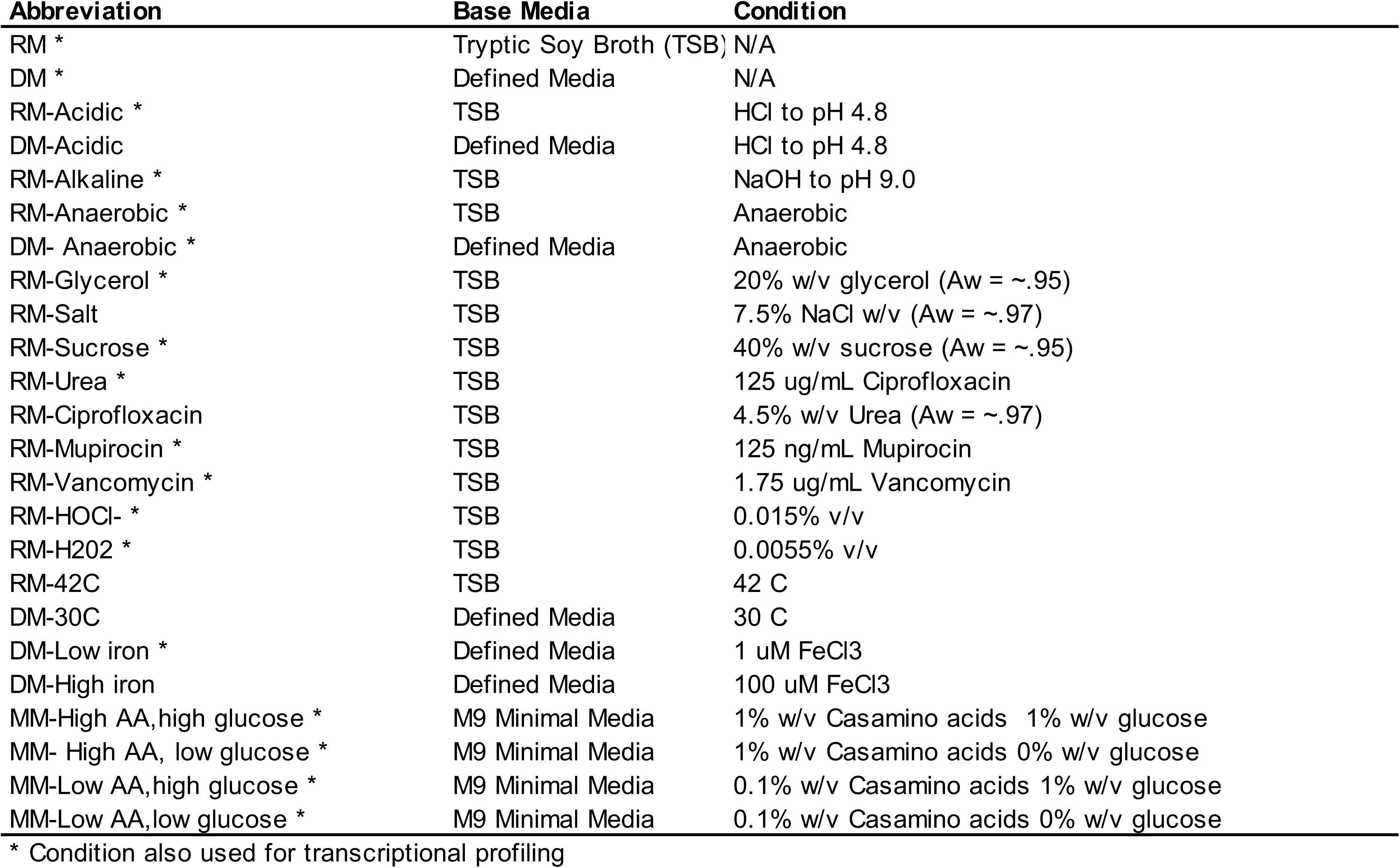
Cultivation Conditions. Gene fitness screening was conducted in 25 diverse conditions, chosen to reflect stressors encountered in colonization and infection. * denotes those conditions where matched samples were also transcriptionally profiled with RNA-seq.

#### Essential gene identification

Due to the varying growth rate of *S. epidermidis* across conditions and the level of growth that each of these conditions supports, we observed varying log2FC distributions across conditions (Fig. 2a). Thus, we decided against a strict log2FC cutoff for gene hits, which would have the effect of over or under-identifying potential ‘hits’, depending on the distribution. Instead, we opted to call hits individually for each condition and then leverage this data across all conditions to determine a set of high-confidence, medium-confidence, and low-confidence putative essential genes (Fig. 2b), identifying 160, 442, and 304 respectively (STable 2-4). To circumvent relying on limited genome annotation and unvalidated operon prediction in *S. epidermidis*, we opted to call hits based on individual genes irrespective of operon organization. As polar effects were estimated to result in a 15% false positive rate in the discovery of essential genes in a CRISPRi screen in *E. coli* (Rousset et al., 2018a), we anticipate that these effects do not represent a major limitation to our study.

**Figure 2.**
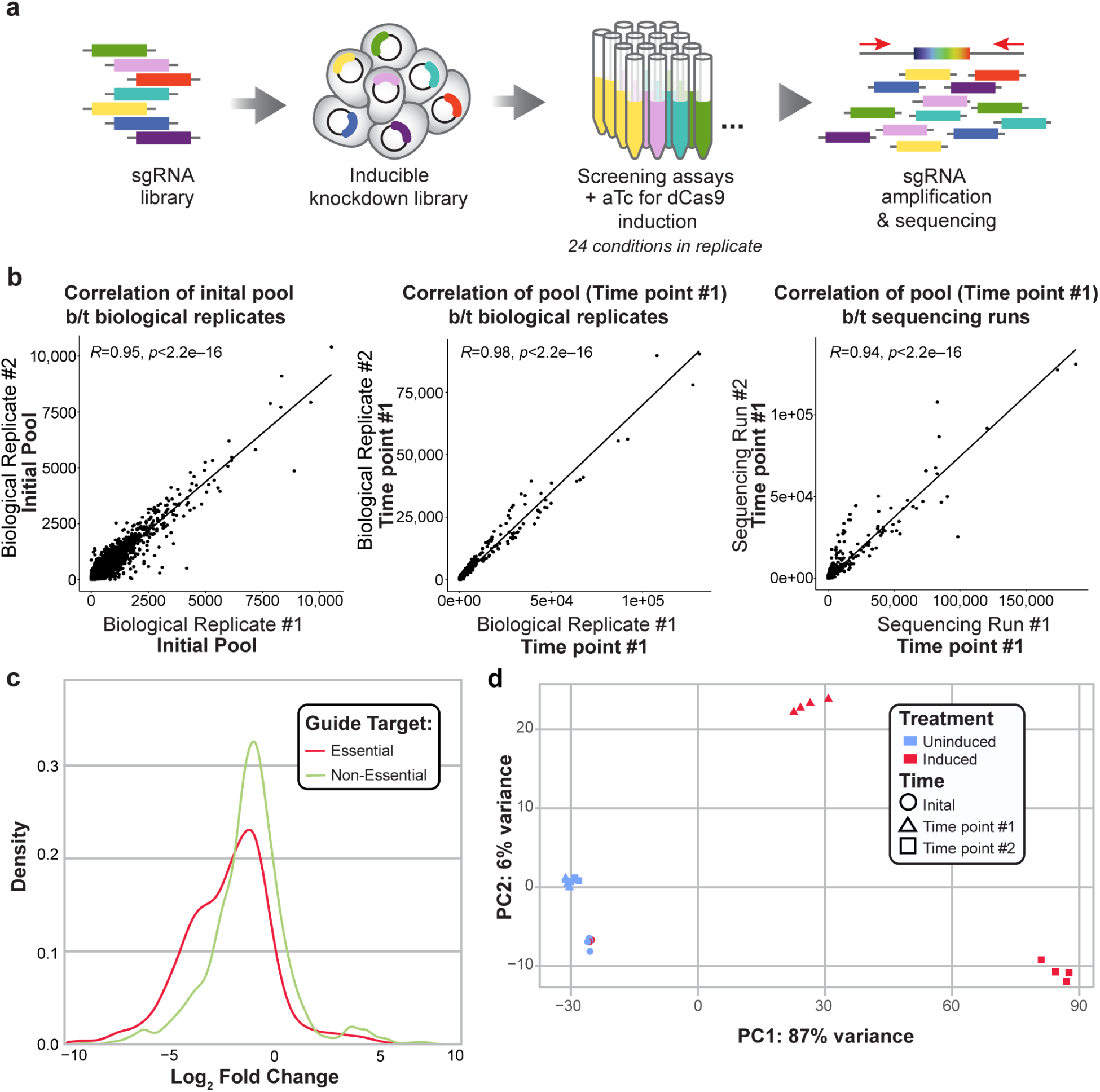
The genome-wide CRISPRi library demonstrates inducible knockdown in rich media (RM). (A) Overview of library creation and testing. Briefly, guides were designed to all available NGG PAM sites in the *S. epidermidis* Tü3298 genome. Guides were synthesized on microarray, ligated into a custom anhydrous tetracycline (aTC)-inducible pidCas9 shuttle vector, and transformed into *S. epidermidis* via sequential transformations into *E. coli*, *S. aureus*, and a final phagemid transfer step. The library was screened using 0.1 uM aTC for inducible knockdown across 25 conditions in at least triplicate and sequenced on Illumina HiSeq and NovaSeq platforms. (B) Biological replicates of normalized read counts of the initial knockdown pool (left panel), after 6 hours of induced growth (center), and between sequencing runs (right) are highly correlated (Pearson correlation, upper left). (C) Density distribution of the Log2 Fold Change (log2FC) of guides targeting putative essential (defined by homology to known *S. aureus* essential genes) and non-essential genes after screening in rich media, where log2FC = relative abundance of a guide in the final pool / initial pool (calculated using DeSeq2). (D) PCA plot of normalized read counts of the initial pool, induced, and uninduced samples after growth to early (Time point #1) and mid-exponential phase (Time point #2). Uninduced samples grown to exponential phase (blue triangles and squares) cluster together near the initial pool (blue circles), which indicates even growth of the control pool. Induced samples grown to exponential phase (red triangles and red squares) cluster together by time point away from the initial pools and uninduced pools, which indicates the impact of induction on the final composition of the pool.

We then examined consistency between conditions as a quality control check and to identify possible technical variation between our diverse conditions. First, we grouped our conditions by background media type – rich, defined, or minimal (Table 1) – and generated a PCA plot of the normalized read counts. As expected, the initial pool samples cluster tightly together and there is slight differentiation between the background media types. Next, we sought to determine the consistency of our screening results across conditions. With the exception of condition-specific essential genes for each stressor or nutrient environment, we expected to see that a gene behaves similarly (e.g., essential or non-essential) across conditions. To determine this, we generated heatmaps of the conditions grouped by media type (median log2FC of all guides for each gene vs. condition). We observed that the median log2FC of each gene was largely consistent between conditions, suggesting little technical variability between screen conditions (SFig. 2a-c).

We then sought to examine the broad functional classifications of these putative essential genes. We performed a KEGG enrichment analysis on the medium and high confidence gene lists, compared to the Tü3298 background, and identified enrichment for pathways involved in genetic information processing, amino acid metabolism, and global metabolism, including aminoacyl-tRNA synthesis, mismatch repair, and homologous recombination (Fig. 2d). Generally, these results were consistent with the established literature regarding gene essentiality in staphylococci (Chaudhuri et al., 2009; Christiansen et al., 2014) and model microorganisms, which have identified DNA and RNA processing and protein synthesis among key essential functions (Gerdes et al., 2003). While previous reports in *S. aureus* demonstrate that genes involved in amino acid metabolism are not consistently identified as essential in rich media (Chaudhuri et al., 2009; Christiansen et al., 2014), amino acid metabolism has been hypothesized to play a role in *S. aureus’* functional adaptation to diverse nutrient environments (Bosi et al., 2016). Our results suggest that amino acid biosynthesis and metabolism are core mechanisms underlying *S. epidermidis’* success across a range of diverse nutrient and stress-related conditions.

Finally, we individually tested this list of high-confidence essential genes to gauge the ability of the pooled assay to identify genes whose knockdown affects growth. We generated ### individual knockdown strains for each of these hits and assessed growth rate in rich media (Fig. 2e). Despite differences afforded by competitive growth in the pooled assay, taken together with sequencing-based quantitation which can identify subtle growth defects, we demonstrated that the majority of these high-confidence hits also demonstrated a growth defect when grown in single culture (vs. a no- knockdown strain).

#### Multi-stress response gene identification

We then asked whether we could use the CRISPRi screen data to identify stress response genes, i.e., those with a fitness defect in multiple stress conditions, which may represent genes responsible for adaptation to stress or other environmental signals. By screening across multiple stressors, we could identify high confidence hits, leveraging the utility of the pooled CRISPRi approach. To control for the effect of background media type (e.g., rich vs. minimal) on gene fitness, we restricted our study of multi-stress response genes to stress screens conducted in rich media (TSB). These conditions included multiple antibiotic stress conditions, multiple hyperosmotic stress conditions, oxidative stress conditions and more (Table 1). We designated a gene as a multi-stress response gene if it was identified in at least ∼50% (6/13) of these rich media stress conditions but not identified as a hit in plain rich media or as a putative essential gene (high or medium confidence). We identified 25 such genes (Table 2), several of which have known functions in stress response or DNA or protein damage repair (e.g., *ssrB, msrA, uvrC, dinG*) (Asad et al., 1995; Dhandayuthapani et al., 2001; Goerlich et al., 1989; Mashruwala and Boyd, 2017; Voloshin et al., 2003; Wilde et al., 2015; and reviewed in Singh et al., 2018) . Of particular interest are previously unidentified hits related to inorganic ion transport and metabolism. We identified three genes in this category (*ptsA*, *cntL*, *cntA*), of which *cntL* and *cntA* are involved in staphylophine biosynthesis and staphylopine-depdendent metal transport, respectively. *PstA* is involved in phosphate and nickel transport, while staphylophine is a broad-spectrum metallophone (Song et al., 2018). Trace metals, including nickel, are vital cofactors for enzymatic catalytic activity, and we postulate that demand for these cofactors is increased under cellular stress. We note that of 25 identified genes, six have no or poor functional annotation. Given their potential role in response to multiple stressors, these genes are strong candidates for future follow-up investigations. Finally, our analysis also identified two genes that are annotated as essential in *S. aureus* (*nusB, menD*). These may be genes that were missed by our essential gene screening, or represent true biological differences in gene fitness between the species.

**Table 2.**
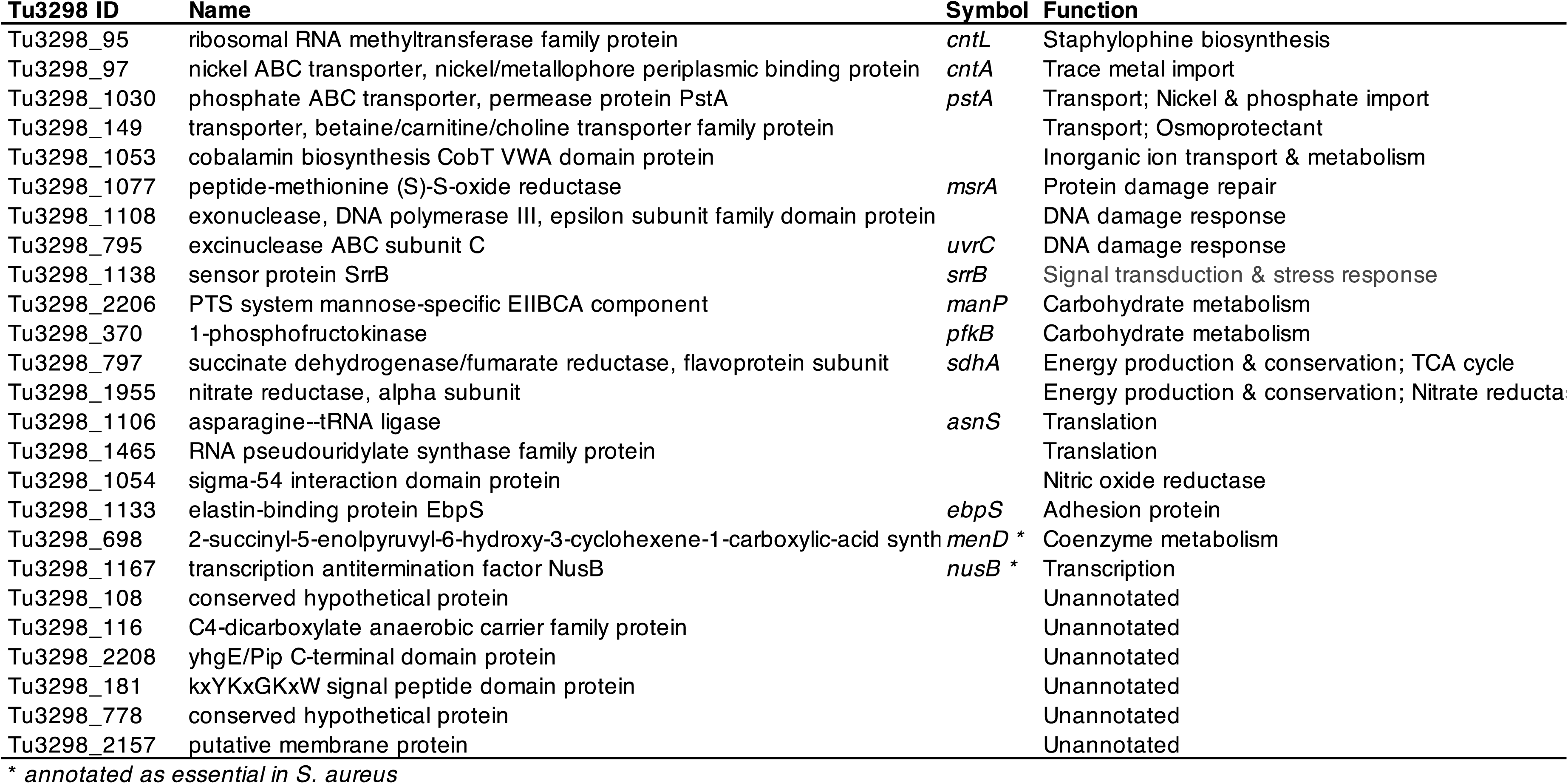
CRISPRi identifies multi-stress response genes in rich media. A gene was designated as a multi-stress response gene if it was identified as a hit by multiple guides in at least 50% of the rich media stress conditions.

Taken together, we demonstrate how high-throughput CRISPRi screens across diverse growth conditions can identify essential gene and multi-stress response gene candidates. With stringent thresholds for gene hit identification – at least 2 independent guide hits per gene and analyses conducted across multiple conditions in at least biological triplicate – these results likely represent robust phenotypes. Our individual confirmation of high-confidence essential genes further supports the validity of such screens.

### CRISPRi screening identifies condition-specific gene hits

We then sought to identify specific genes vital to growth in each condition, which would provide insights into *S. epidermidis’* lifestyle versatility. By identifying hits specific to each environment, we could also construct an accessible resource of gene fitness data for this difficult-to-study microbe. Here, we considered condition-specific genes as hits identified under a particular condition, but not considered a high or medium confidence putative essential gene (i.e., those identified across all conditions; full results shown in Supplemental Tables 5-28). We note an inherent limitation of our approach is that our library does not cover every protein coding sequence and, as such, the absence of a gene as a hit should not be taken as evidence that the gene is *not* conditionally essential. Rather, this condition-specific essentiality screen provides high quality hits that constitute a starting point for investigating genes of interest.

#### CRISPRi screening suggests a role for urease and cell wall function in response to salt stress

As an example, we asked whether we could identify genes required for survival under high salt stress, a stress condition that *S. epidermidis* may face on the surface of human skin. After performing the fitness assay screening in 7.5% sodium chloride, we identified several condition-specific hits previously identified in the literature as vital under high salt stress, including a cation transporter, genes of the *dlt* operon (implicated in cell wall biosynthesis), and a beta/carnitine/choline transporter (Kappes et al., 1996). We also found several previously unidentified hits, including two urease accessory coenzymes involved in nickel binding (UreG, UreE), a transcriptional regulator annotated only as involved in “cell envelope-related function” (Tü3298_708), and a teichoic acid D-alanine hydrolase (*fmtA*). Although the exact function of *fmtA* is unknown, it is implicated in cell wall structure and methicillin resistance in *S. aureus* (Komatsuzawa et al., 1999). Given the role of cell wall structure in response to hyperosmotic stress, we speculate that this is the basis for the essentiality of *fmtA* and the putative transcriptional regulator gene in the high salt stress condition. The role of the accessory urease chaperones in high salt stress is a curious find and, taken together with other high-quality hits (*secA2*, *purH*, *purM*), are of interest for follow-up investigation into the role of salt tolerance in *S. epidermidis*.

### RNA-seq identifies pathways involved in *S. epidermidis* stress response and growth in diverse nutrient conditions

#### Overview of transcriptomics analysis across conditions

While fitness investigations identify genes necessary for growth in stress conditions or under nutrient limitation, RNA sequencing studies define the global transcriptional plasticity of an organism. Indeed, it is well established that differential expression of a given gene is a poor indicator of gene fitness under stress (i.e., many differentially expressed genes are not essential), and thus transcriptomic analysis provides a complementary approach to studying lifestyle versatility in *S. epidermidis*. A subset of 18 conditions from the CRISPRi screening assays, selected to represent a diversity of stressors across minimal, defined, and rich background media, was selected for RNA-seq to examine pathways involved in *S. epidermidis* stress response to growth in diverse nutrient conditions. To broadly examine *S. epidermidis* adaptation across conditions, we first grouped conditions by background media type, as we anticipated that nutrient availability would have the strongest influence on transcription, confirmed by PCA analysis (Fig. 4a).

**Figure 3.**
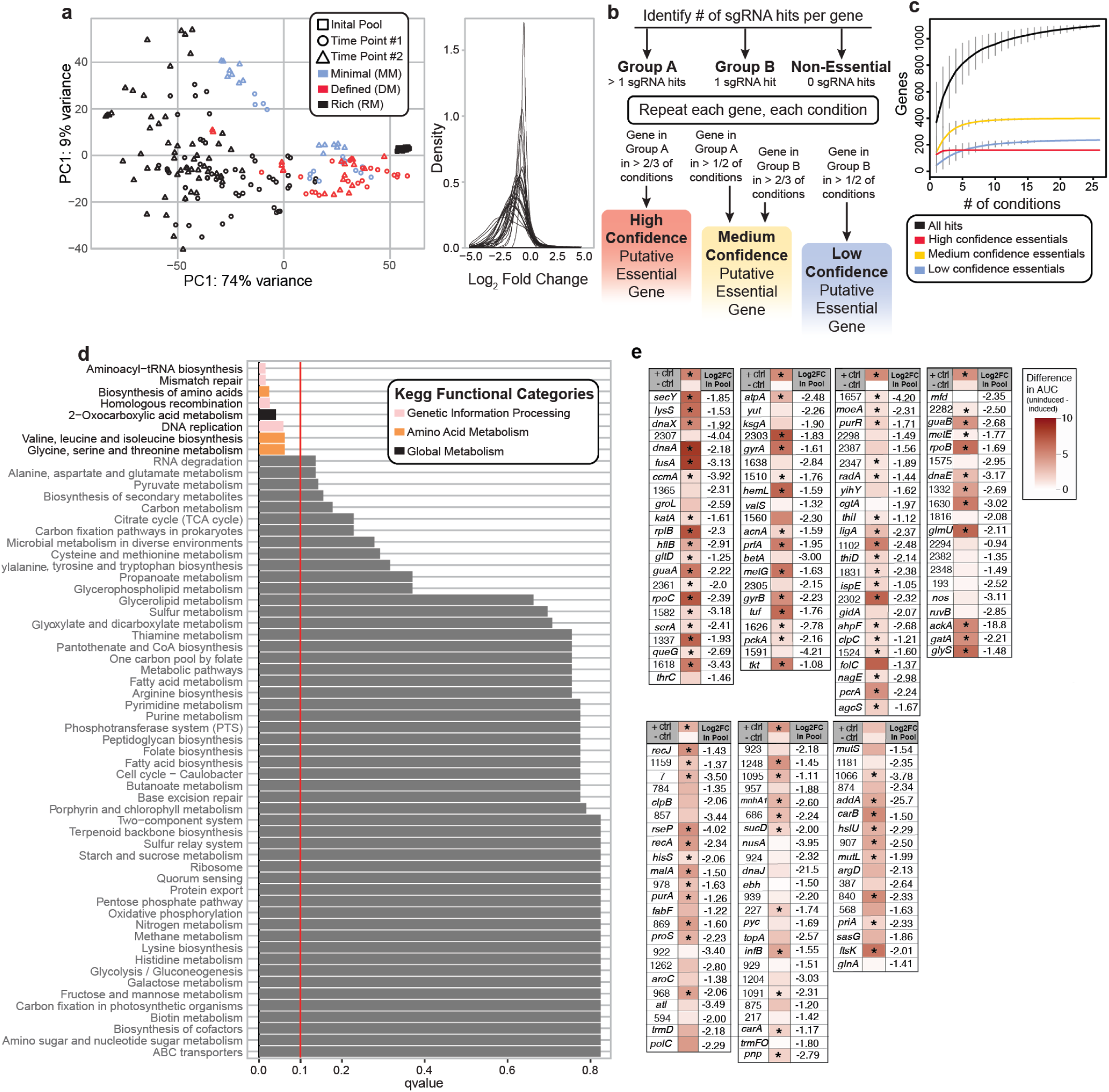
Multi-condition CRISPRi identifies putative essential genes. (A) PCA plot of normalized read counts demonstrating segregation of samples by media type and time point after aTC induction (initial pool is also plotted, clustered as rightmost overlapping black squares). (B) Workflow of putative essential gene identification across conditions. Identified genes were binned into high, medium, or low confidence categories based on quality of supporting data as described. (C) Gene accumulation curves for high, medium, and low confidence putative essential or non-essential genes across increasing numbers of conditions. (D) Enriched KEGG pathways within the high and medium-confidence essential gene list. Significant pathways (as determined by default GAGE analysis (parametric t-test and multiple test correction to control FDR); FDR < 0.10) are colored by KEGG category; gray pathways indicate non-significance. Vertical red line indicates our significance threshold of FDR = 0.10. (E) Validation of high confidence hits. Difference in area under growth curve (AUC): uninduced AUC (i.e., no knockdown) – induced (i.e., knockdown) AUC depicted in red. Significance testing: Student’s t-test for AUC uninduced vs induced; n = 3; * = pvalue < 0.05. + ctrl is known essential gene Tü3298_1803 (ribosomal protein L2) - ctrl is known non-essential gene Tü3298_1121(nucleoside diphosphate kinase). P-value for +ctrl in last batch = 0.055. Gene name provided if annotated, otherwise. #= Tu3298 Gene ID

**Figure 4.**
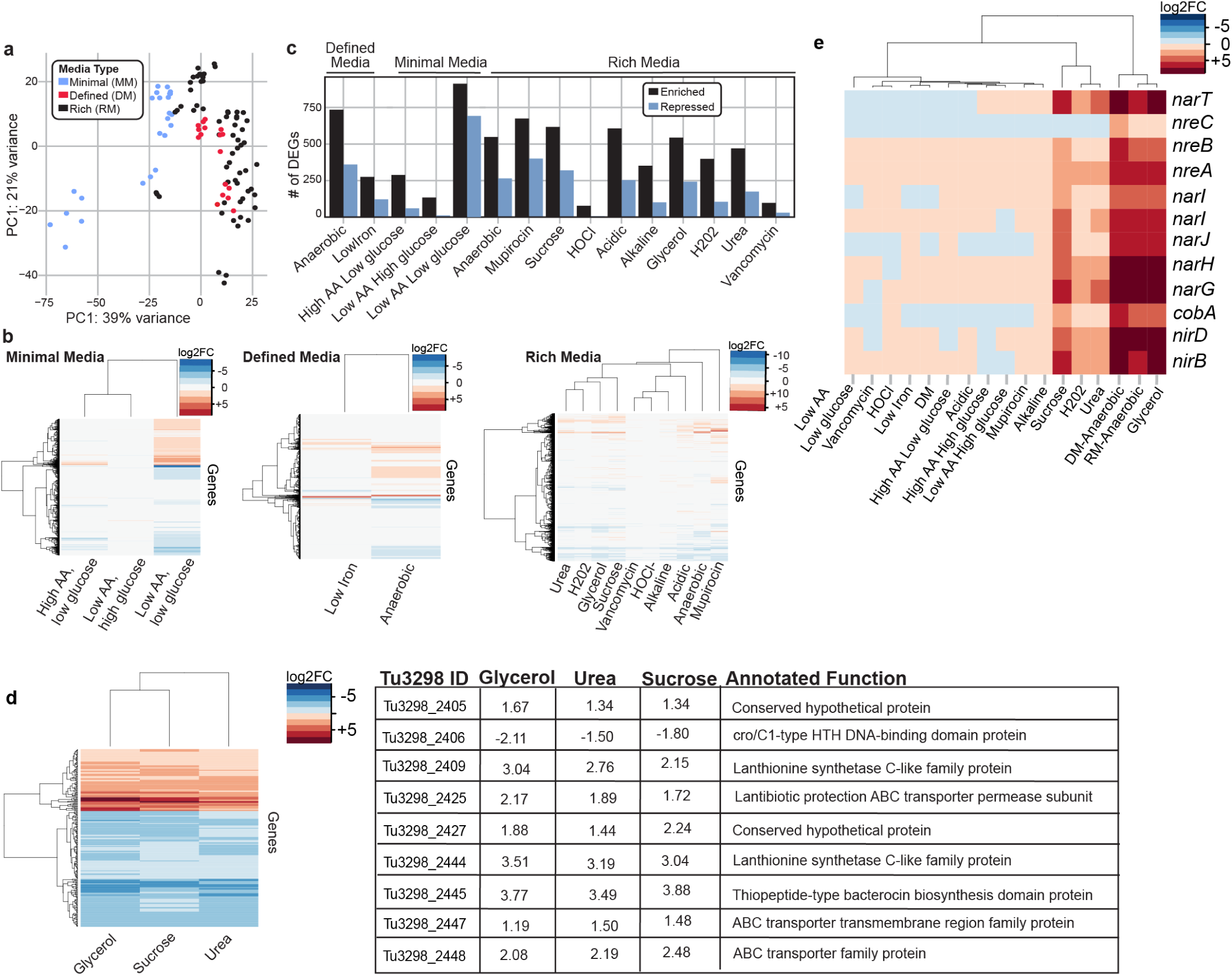
Differentially Expressed Genes (DEG)s are identified by matched transcriptomic analysis of CRISPRi growth conditions. (A) PCA plot of normalized read counts depicting clustering of RNA-seq samples (n=3 per condition) by media type. (B) Heatmaps of log2FC for each condition, grouped by media type. Reference for RM conditions = plain RM (TSB), DM conditions = plain DM, MM conditions = M9 MM supplemented with 1.0% CAA and 1.0% glucose (high amino acid, high glucose condition). (C) Number of DEGs for each condition, grouped by media type. (D) Heatmap of genes up (log2FC >1) or down (log2FC <-1) regulated in all of the osmotic stress conditions. Table of genes involved in lantibiotic synthesis and export differentially expressed in all three osmotic stress conditions. (E) Heatmap of genes involved in nitrate/nitrite reduction across conditions as compared to RM. Hierarchical clustering for (B), (D), (E) was based on Euclidean distances.

#### Global transcriptomic analysis identifies genes enriched in hyperosmotic stress conditions

One advantage to performing transcriptomic analysis across a variety of stress conditions is the ability to identify overlapping responses between conditions within a stressor type (e.g., osmotic stress, antibiotic stress, oxygen limitation) to further inform cellular response. As an example, we sought to identify a subset of genes globally responsive to hyperosmotic stress. We identified 209 genes significantly upregulated in all three hyperosmotic stress conditions (sucrose, glycerol, urea) with a log2FC cutoff of 0.5 and 81 genes with a more stringent log2FC of 1.0 (Supplemental Tables 54 & 55). KEGG pathway analysis on the most highly enriched genes (log2FC > 1) identified the following categories as significantly enriched (FDR < 0.1) under osmotic stress: nitrogen metabolism, two-component systems, butanoate metabolism, microbial metabolism in diverse environments, glycolysis/gluconeogenesis, purine metabolism, and propanoate metabolism. Consistent with previous reports, we identified a choline-glycine betaine transporter (Tü3298_149) and genes involved in iron uptake (*feoA, feoB*)(Bojanovič et al., 2017; Hengge-Aronis, 1996; Ly et al., 2004). Interestingly, multiple genes involved in bacteriocin synthesis were enriched (highlighted in Fig. 4e). Specifically, gene clusters involved in epidermin synthesis and export were enriched across all three osmotic stress conditions. Epidermin is a pore-forming antimicrobial peptide effective against a range of Gram-positive bacteria, including *S. aureus*. While the role of bacteriocin synthesis and export during osmotic stress is not readily apparent, one possibility is that it may be a generalized mechanism to improve competition with other members of the skin microbiome during periods of stress.

#### Global transcriptomic analysis identifies nitrogen reduction as enriched in anaerobic, hyperosmotic, and hydrogen peroxide stress

We were also surprised to find genes involved in nitrogen metabolism as some of the most highly enriched in all three hyperosmotic stress conditions. We saw significant enrichment of genes involved in nitrate/nitrite transport (*narT*), oxygen regulation of nitrate/nitrite reduction (*nreA, nreB*), nitrate reduction (*nar* operon), and nitrite reduction (*nir* operon) (Fig. 4d). While a few studies have suggested overexpression of the nitrate or nitrite reductase complexes during osmotic stress, no clear mechanism has been elucidated(Finn et al., 2015; Withman et al., 2013). In a study of uropathogenic *E. coli*, Withman et al. speculated that by linking hyperosmotic gene expression to anaerobic gene expression (e.g., overexpression of nitrite/nitrate reductases), *E. coli* may adapt more readily to the hyperosmotic and hypoxic environment of the urethra and bladder lumen. Similarly, *S. epidermidis* must combat hyperosmotic stress on the skin, which results as salt and urea-rich sweat evaporates from the surface. We speculate that the upregulation of nitrate/nitrite reduction genes in response to hyperosmotic stress provides an avenue for *S. epidermidis* to utilize nitrogen from sweat.

Given this striking finding, we then asked whether these genes were similarly enriched in any other stress conditions compared to rich media. Enrichment of nitrate/nitrite reduction genes is expected under anaerobic conditions, where nitrate is used as an electron acceptor during respiration and, indeed, these genes are enriched under both anaerobic defined and rich media. In addition to enrichment under hyperosmotic stress, nitrate/nitrite reduction genes were significantly upregulated in response to hydrogen peroxide stress (Fig. 4d). Under this condition we also identified upregulation of additional anaerobic metabolic genes, including *pflB* (log2FC = 4.29) and *nrdG* (log2FC = 2.12), among others. *S. aureus* has been shown to upregulate genes involved in anaerobic metabolism in response to hydrogen peroxide induced oxidative stress, indicating that the microbe undergoes an oxygen-limiting state after exposure (Chang et al., 2006). Our findings here suggest a similar mechanism for *S. epidermidis* response to hydrogen peroxide stress.

#### Condition-specific RNA-sequencing generated a rich resource for investigating the transcriptomic response of S. epidermidis across a diversity of conditions

Next, we sought to investigate differentially expressed genes (DEGs) within each condition. We expected that by analyzing DEGs (and corresponding KEGG pathways), we could investigate the overall transcriptomic response of *S. epidermidis* in a variety of environmental conditions (Fig. 5) We identified differentially expressed genes (log2FC > 0.5 or < -0.5 and FDR < 0.1) in each condition compared to their background media type (see Supplemental Tables 30-53 for all log2FC data for each condition, including FDR-adjusted p-values values). We found that generally, stress and nutrient limited conditions resulted in significantly more upregulated genes/corresponding KEGG pathways than repressed genes (Fig. 4c), likely reflecting a broad transcriptional stress response, or large-scale mobilization of genes to increase resource utilization.

**Figure 5.**
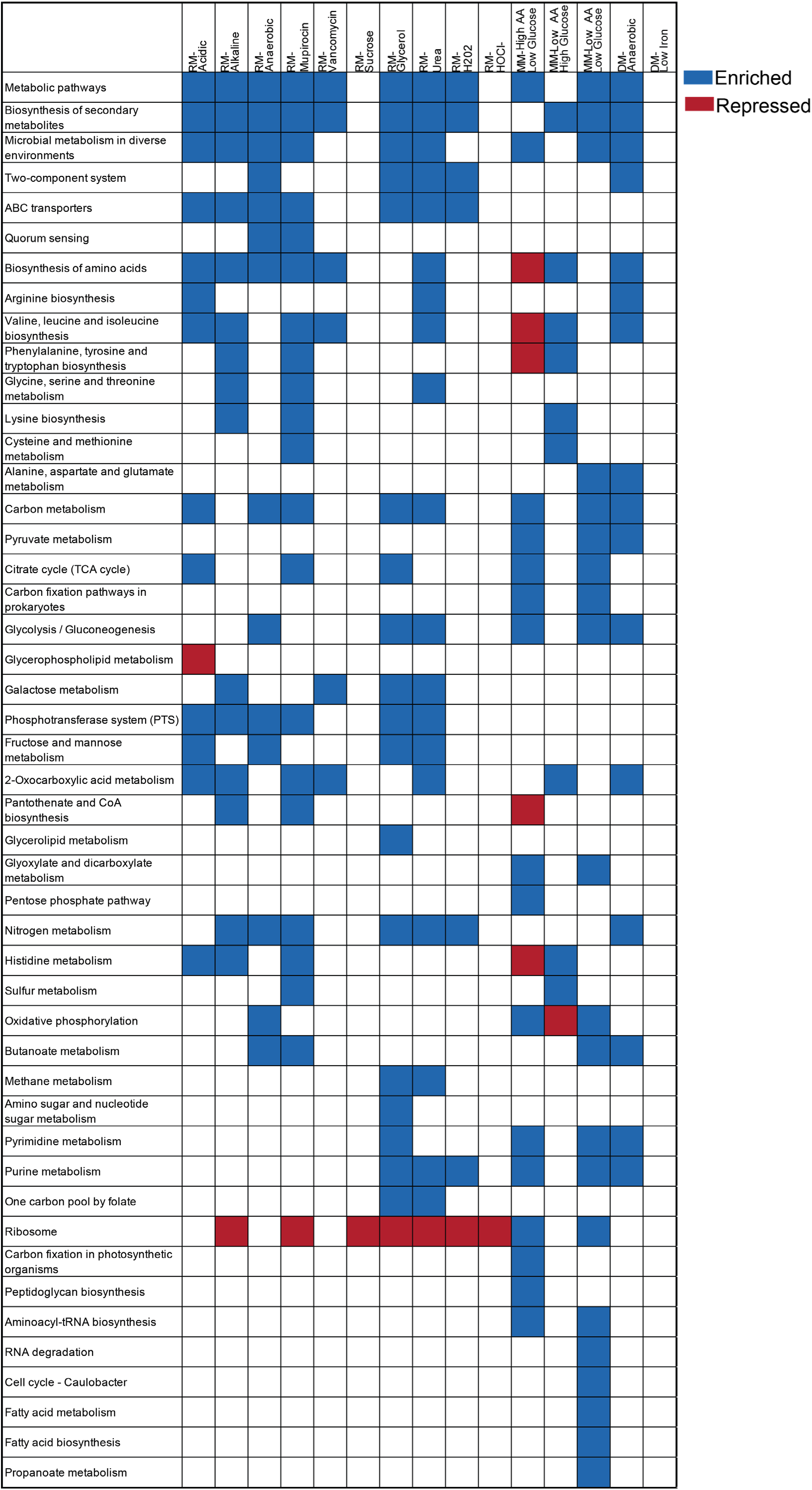
RNA-seq identifies pathways involved in S*. epidermidis* plasticity across diverse environments. The top 5 KEGG pathways most significantly enriched and repressed in each condition are grouped by media type. RM samples are compared to RM baseline media, DM samples are compared to DM baseline media and MM samples are compared to high amino acid (1.0% CAA), high glucose (1.0%) minimal media. (+) represents significantly upregulated KEGG pathways at FDR < 0.10; (-) represent significantly downregulated pathways at FDR < 0.10. For all KEGG pathways significantly up and down regulated for each condition, see Supplemental Table 56.

**Figure 6.**
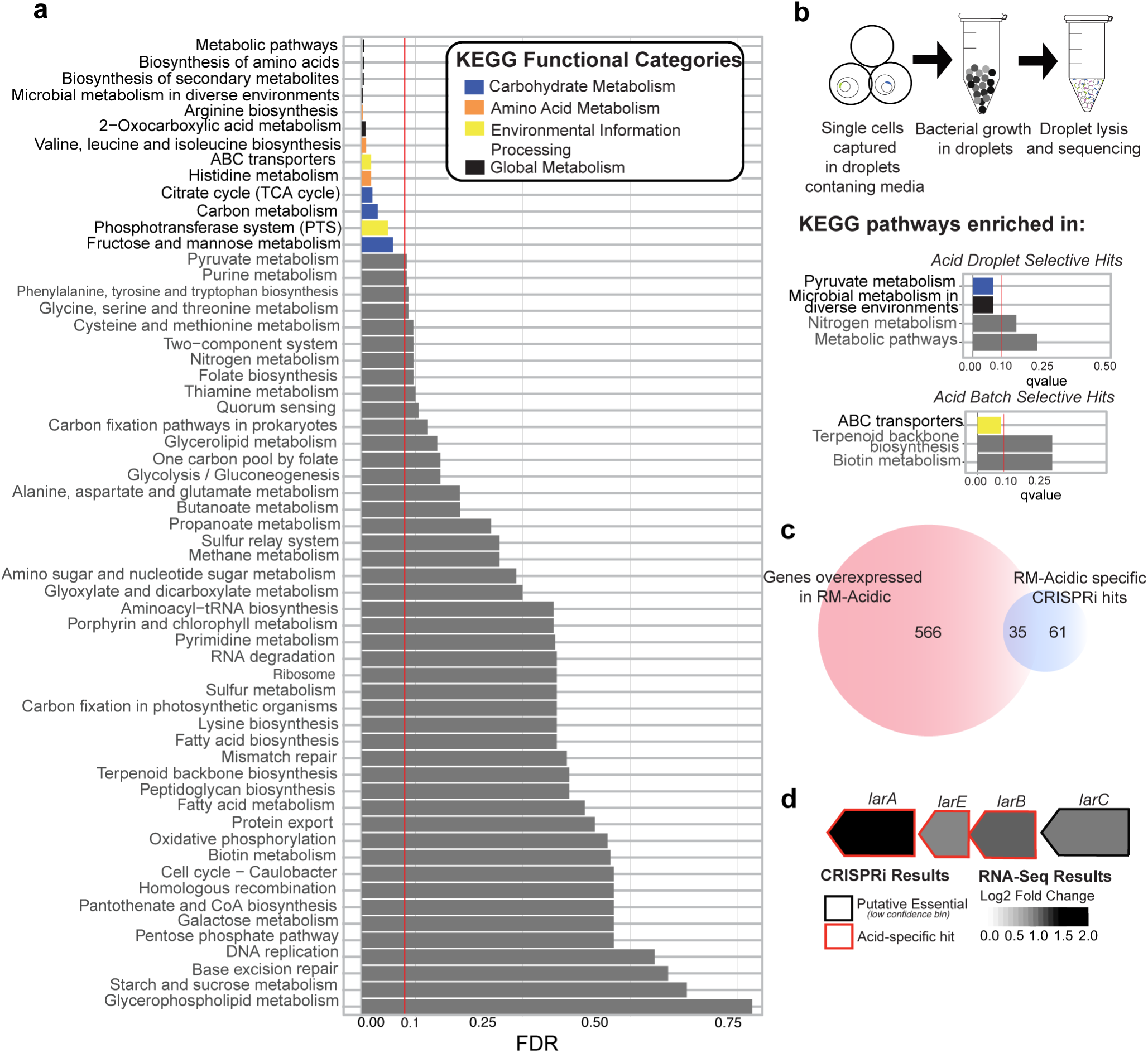
Integrated gene fitness screening and transcriptomics analysis identify cellular functions necessary for survival and adaptability under acid stress. (A) KEGG pathways enriched in RM-acidic media (vs. RM) by RNA-seq. Statistically significant results are colored by KEGG category; grey bars represent non-significant results. Vertical red line indicates our significance threshold of FDR = 0.10. (B) Workflow for droplet-based CRISPRi screening. Briefly, single cells captured in droplets containing RM-acidic media are incubated to promote growth. After incubation, droplets are lysed, and the lysate is amplified and sequenced as for a batch pool. KEGG pathways enriched in droplet-specific and batch-specific gene lists. Pathways with FDR ≤ 0.25 shown. Significantly enriched pathways (FDR ≤ 0.1) are colored by KEGG category. (C) Venn diagram depicting overlap of genes overexpressed in acidic media and genes identified as acid-stress specific hits by CRISPRi. (D) Lactate racemization operon (*lar)* with log2FC (RM-acidic vs. RM) and CRISPRi screening results for each gene.

As an example, we investigated the transcriptional response to growth in an anaerobic environment. DEGs and KEGG pathways enriched under anaerobic conditions were generally consistent with previous reports. For example, the most highly upregulated genes were involved in nitrate reductase (*nir* and *nar* operons) and oxygen sensing (*nreB, nreC*) (Fedtke et al., 2002; Schlag et al., 2008). Genes of the urease operon were also among those most highly upregulated during anaerobic growth, perhaps as a response to an altered pH in the anaerobic condition. Interestingly, three genes of the beta class of phenol soluble modulins (PSMβ) were highly enriched in the anaerobic condition (log2FC = 5.91, 5.77, & 5.53), along with the expression of the quorum sensing *agr* locus. Literature regarding the expression of the quorum sensing system in hypoxia is conflicting; in our strain of *S. epidermidis* at the least, our results provide evidence for upregulation of quorum sensing genes and downstream effectors under oxygen limitation. We speculate that regulation of the quorum sensing system, which controls virulence and other metabolic processes in staphylococci (Batzilla et al., 2006; Cheung et al., 2014; Queck et al., 2008), is one mechanism by which *S. epidermidis* adapts to the differential oxygen levels of infection niches.

### CRISPRi screening and RNA sequencing cooperatively identifies pathways vital for acid resistance in *S. epidermidis*

While CRISPRi and transcriptomics each are powerful tools for probing gene function, integrating these datatypes could provide complementary data on potential gene function. As an example, we performed such an analysis on the acid stress condition, which is relevant to the widely varying pH in human skin (4.1-5.8, depending on skin site, individual, and the lifestyle factors (Lambers et al., 2006; Segger et al., 2008)). While several known strategies have been identified in bacteria to accommodate changes in environmental pH, including proton gradients, generation of alkaline products (e.g., ammonium production, amino acid neutralization pathways), alterations to the cell membrane, metabolic changes, and more (reviewed in (Guan and Liu, 2020)), little is known about the acid response system in *S. epidermidis.* In addition to examining transcriptomic and gene fitness data, we also piloted a droplet-based CRISPRi approach to identify growth defects in genes that may be masked by the effects of growth in batch culture.

#### Droplet-based CRISPRi identifies distinct, conditionally essential genes compared to batch CRISPRi

One disadvantage of batch culture CRISPRi as performed is the inability to identify growth defects in knockdown strains that may be masked due to metabolic cross-feeding or the secretion of other public goods. To examine what new insights could be gained by circumventing this limitation, we performed a droplet-based CRISPRi in acid stress. Here, individual knockdown strains are captured in droplets containing growth media, grown individually, and then pooled for sequencing for quantitation. While we noted a technical limitation in the lower diversity of the knockdown library recovered after droplet-based growth (likely due to incomplete capture of knockdown strains in droplets), we identified a distinct set of genes vital for growth in acidic media.

Consistent with our batch CRISPRi screening, condition-specific hits here are those that are not identified as a putative essential gene (medium or high confidence). Additionally, to help control for differences in gene fitness due to droplet growth, genes were only deemed an acid-specific droplet ‘hit’ if they were not also identified in a droplet-based rich media screen. 50 genes were identified as acid-specific hits by at least two independent guides (STable 56), including genes involved in transport, energy production and conservation, or cell wall biogenesis; 14 of the 50 hits had no or poor functional annotation. Several of our identified hits in the droplet-based assay have been previously implicated in response to acid stress, including: nitrate/nitrite reductases (Moran et al., 2017), multiple amino acid permeases, and a Na+/H+ transporter (reviewed in (Guan and Liu, 2020)). Comparatively, 96 genes were identified as acid-specific hits by at least two independent guides in batch growth (STable 57), including ABC transporters and genes involved in coenzyme metabolism, cell wall biogenesis, energy production and conservation, or amino acid metabolism (among others), with 21 hits having no or poor functional annotation. As with our droplet-based assay hits, many of these batch hits have been previously identified in response to acid stress, including: multiple amino acid permeases, arginine kinase (Canonaco et al., 2003), and carbamate kinase (*arcC*) (Casiano-Colón and Marquis, 1988). Surprisingly, only 17 genes were identified by both approaches, including transporters (e.g., a Na+/H+ antiporter), genes involved in cell wall biogenesis, and four genes with poor/no annotation (Table 3). Though we found the two datasets sparsely overlapping, these 17 genes are likely high confidence hits for validation.

**Table 3.**
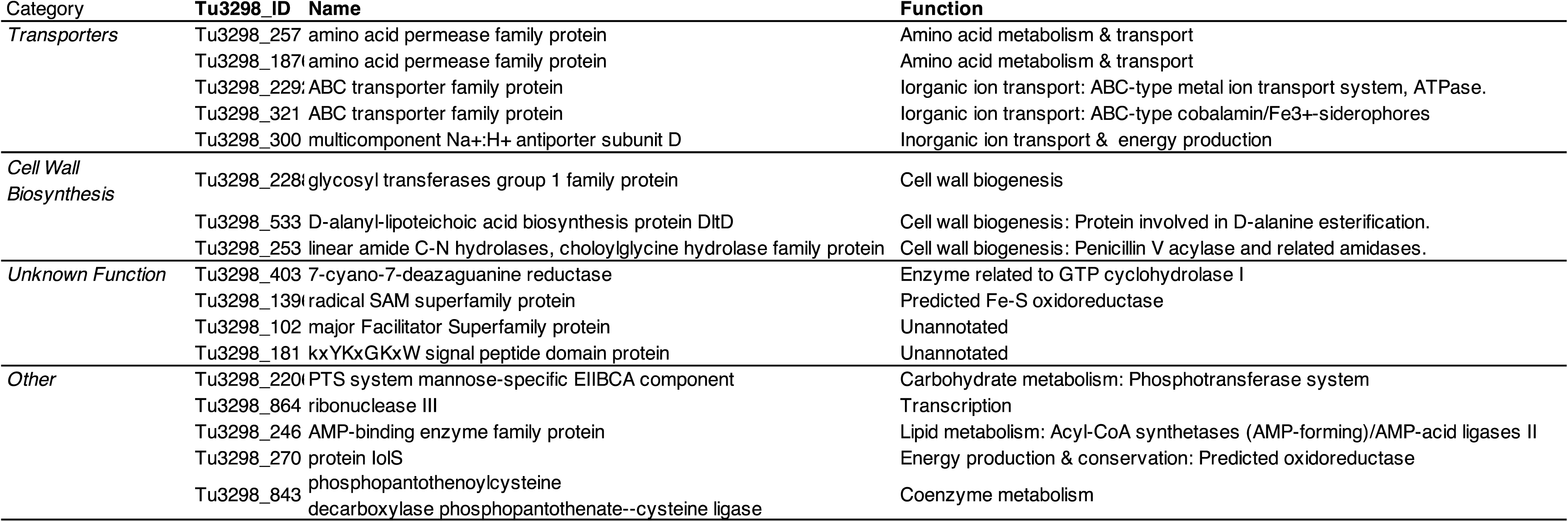
Genes identified by both batch and droplet-based CRISPRi. . 17 genes were identified as a hit by two or more guides in both the batch-based and droplet-based CRISPRi screens, indicating a high-confidence hit.

Finally, we examined gene hits unique to a method (i.e., “droplet selective” vs. “batch selective” hits). A KEGG enrichment analysis identified significant enrichment of “pyruvate metabolism” and “microbial metabolism in diverse environments” for droplet-selective hits, and “ABC transporters” in batch-selective hits (Fig. 5c). In addition to these enriched KEGG categories, we also identified individual genes that are exclusive to droplet-based or batch screening, such as the nitrate/nitrite reductase genes seen exclusively in our droplet screen. We speculate that disruptions in pyruvate metabolism lead to altered fermentative byproduct production in the confined droplet space, where there is less diffusion, with the corollary that increased accumulation of metabolic byproducts may lead to a lower tolerance to existing acid stress. Similarly, we postulate that the identification of the nitrate/nitrite reduction systems exclusively in the droplet-based screen may be related to ammonium production and export in the confined droplet condition. Our basis for this hypothesis includes previous data indicating that nitrate/nitrite reductase genes are upregulated in response to acid stress in *S. epidermidis* and are thought to contribute to acid tolerance through the production of basic ammonia (Moran et al., 2017). In addition, some microbial species combat acid stress through secretion of this excess ammonia, which raises the pH of the immediate extracellular environment (Vylkova, 2017). Interestingly, we observed a significant clumping phenotype in this condition, suggesting that cells preferentially grow in close proximity (vs. complete planktonic growth). We speculate that in batch CRISPRi screening, nitrate/nitrite reductase knockdown strains may not demonstrate a fitness defect if they grow in close proximity to other strains that are producing ammonium (which would raise the localized pH or be available for import by deficient strains). This is in contrast to the homogeneity of droplets, in which cells growing in close proximity are composed of a single knockdown strain (SFig 3a-c) and are not thereby influenced by extracellular products of different knockdown strains. Though speculative, these proposed mechanistic links could partially explain the distinct hits that we observe in the droplet-based assay (nevertheless, we note that technical differences between the two approaches also likely have some effect). Overall, our suggest that there is a utility for both the batch and droplet-based approach in identifying genes of interest using CRISPRi, although future investigations should focus on improving technical limitations of the droplet system (e.g., lowered library diversity as a result of incomplete capture).

#### RNA-seq identifies genes enhanced and repressed in acidic media

We then complemented gene fitness data with transcriptomic analysis in the same acid environment. KEGG pathway analysis identified significant enrichment of pathways involved in amino acid metabolism, carbohydrate metabolism and environmental information processing (Fig. 5a). Specifically, significantly upregulated pathways by Generally Applicable Gene-set Enrichment (GAGE) analysis include: 1) arginine biosynthesis, 2) microbial metabolism in diverse environments, 3) biosynthesis of amino acids, 4) 2-oxocarboxylic acid metabolism, 5) valine, leucine and isoleucine biosynthesis, 6) ABC transporters, 7) histidine metabolism, 8) tricarboxylic acid (TCA) cycle, and 9) carbon metabolism. Overall, these results are consistent with previous studies in *S. aureus* which suggest a role for amino acid metabolism (particularly arginine), transport and TCA cycle alterations in acid tolerance (reviewed in (Zhou and Fey, 2020)). Among the mostly highly upregulated genes (log2FC >2), we identified efflux pumps, a H+/Na+ antiporter, and components of the arginine deiminase system, which have a demonstrated role in responding to acid stress (Shabayek and Spellerberg, 2017). Surprisingly, however, four genes of the *potRABCD* operon, which codes for a spermidine / putrescene uptake system, were highly upregulated (log2FC *potR* = 2.0, *potB* = 2.1, *potC* = 2.08, *potA =* 1.60). *speG*, which codes for a spermidine acetyltransferase, and *speC*, which codes for an ornithine decarboxylase, were also modestly upregulated under acid stress (log2FC *speG*= 0.557, *speC* = 0.343). While spermidine and putrescene protect against stress in *E. coli* via their role in the glutamate decarboxylase acid response system (Chattopadhyay and Tabor, 2013), this system does not appear to be present our strain of *S. epidermidis*.

#### Leveraging both CRISPRi and RNA-seq analysis identifies a subset of genes enriched and essential in response to acid stress

Previous reports have shown that condition-specific essential genes are not necessarily overexpressed in that condition, possibly to protect vital genes from large fluctuations in transcription (Jensen et al., 2017; Rousset et al., 2021). However, by identifying subsets of genes whose expression is elevated, and who are essential for growth in a stressor, we can begin to conceptually link how a strain’s transcriptomic response is linked to survival under stress. In acid stress, we identified 35 genes both enriched and essential under acid stress (Fig. 5d). Many of these are involved in transport (10 of 35) or cell wall biosynthesis (5 of 35). Of particular interest was the *lar* operon, (Fig. 5d), which encodes a nickel-dependent lactate racemase and several genes involved in the processing and maturation of the enzyme. The role of this lactate racemase is not fully elucidated beyond its role in lactate-consuming microbes. For example, evidence in *Lactobacilllus planatarum* demonstrated that lactate racemase plays a role in cell wall biosynthesis through its generation of D-lactate from L-lactate, consequently conferring resistance to high levels of vancomycin. As previously discussed, modification of the cell wall has been shown to play a role in response to acid stress in *E.coli* (Xu et al., 2020). Taken together, we speculate that the importance of the lactate racemase operon in acid stress in *S. epidermidis* may similarly be related to cell wall structure.

## Discussion

Until the advent of CRISPRi, large scale studies of gene function have been largely limited to genetically tractable model organisms. Circumventing direct gene editing, highly customizable to a genome of interest, and easily scalable, CRISPRi is well suited to study non model microorganisms such as *S. epidermidis,* which despite its ubiquity and unquestionable human heath relevance, has not seen as deep a probe of its function as needed to begin to understand its lifestyle versatility. Here, we present a high-throughput demonstration of CRISPRi in a non-model microbe, complemented with transcriptomics across a diverse array of conditions. Additionally, we piloted an application of high-throughput droplet-based CRISPRi to study gene function in any microbe. Our findings provide a reference both for other staphylococcal researchers and also the prokaryotic community more generally, with our benchmarking analysis for large-scale CRISPRi in a non-model microorganism, which suggests that the behavior of CRISPRi screens are likely species and/or strain-specific (Supplemental Note).

As both a colonizer and pathogen that thrives across human skin and multiple infection niches, *S. epidermidis* adapts to a wide range of environmental stressors. Our fitness screen across multiple conditions in *S. epidermidis* allowed us to identify putative essential genes, multi-stress response genes, and condition-specific essentials. These data then allowed us to make numerous new hypotheses into *S. epidermidis’* biology.

We first investigated this lifestyle versatility at the gene fitness level to identify genes crucial to survival during colonization and infection. In contrast to traditional gene essentiality screens conducted in rich media, we leveraged screen data across diverse environmental conditions and identified genes and pathways, such as amino acid metabolism, that were more broadly vital for growth and adaptability to multiple environmental pressures. For example, we identified staphylophine biosynthesis and dependent metal transport as a crucial stress-response function. Future investigations might focus on the role of staphylophine in colonization and infection niches of *S. epidermidis*; is staphylophine dispensable for colonization of healthy skin but crucial to combat infection-associated stress? Similar investigations could focus on any one of these 26 stress-responsive genes, particularly those with no/poorly annotated function, to determine their importance in healthy colonization versus infection.

Additionally. we complemented our CRISPRi screen with RNA-sequencing to investigate lifestyle flexibility by studying both fitness and transcriptional plasticity. Interestingly, we identified upregulation of the quorum sensing *agr* locus and three PSMβ genes in the rich anaerobic growth condition, suggesting a role for these virulence determinants in a hypoxic infection niche.

Additionally, we observed a surprising enrichment of bacteriocin synthesis/export and nitrate/nitrite reductase genes across three hyperosmotic stress environments, a proxy for the hyperosmotic conditions of sweat. We postulate that nitrate/nitrite reductase upregulation is an adaptation to utilize nitrogen from urea-rich sweat as an energy source, highlighting the role of transcriptional plasticity and an example of the decoupling of gene essentiality and environmental adaptation, as these genes were not crucial for its survival in this environment. This interplay between nitrogen metabolism and hyperosmotic stress adaptation is another potential link for follow-up mechanistic studies.

*S. epidermidis* must combat varying pH levels on skin, which generally ranges from ∼4.1-5.8 but is affected by skin site, individual, and lifestyle habits (Lambers et al., 2006; Segger et al., 2008). Thus, acid stress adaptation is an important component of *S. epidermidis* colonization and lifestyle flexibility on human skin. Our results led us to some intriguing hypotheses into *S. epidermidis’* response to acid stress: 1) polyamine accumulation via an GADR-independent pathway. Polyamines are known for their role in stress response (particularly oxidative stress) across multiple organisms, but the literature regarding their function in acid stress tolerance is sparse. Chattopadhyay and Tabor et al. demonstrated that polyamines protect against aid stress in *E. coli* (Chattopadhyay and Tabor, 2013), but proposed a mechanism via the GADR system, which has no homolog in our strain of *S. epidermidis*; 2) ammonium production by nitrate/nitrite to raise the localized pH, which we only identified by droplet-based CRISPR; and 3) cell wall synthesis and modification, especially the lactate racemase (*lar)* operon, which has previously been demonstrated as important in acid tolerance in *E. coli? S. aureus?* (Xu et al., 2020).

As with any knockdown screening method, inherent technical considerations of CRISPRi resulting in incomplete or ineffective knockdown limit our ability to draw fully conclusive results. First, we acknowledge that we did not target all predicted coding sequences. To the best of our efforts, which involved > 100 transformations into *E. coli* and > 100 subsequent transformations into the intermediary *S. aureus,* we found diminishing returns in efforts invested in guide recovery, as guides are lost, presumably at random, at each step. In addition, 84 coding sequences do not have any NGG targets. Future efforts could supplement pools with sub-pools of synthesized guides to improve genome coverage, or use PAM-less dCas proteins to improve guide coverage across the genome (Chatterjee et al., 2020; Walton et al., 2020).

Second, inherent technical considerations that may limit the ability of the screen to fully identify essential genes, including transcript stability, transcript and protein half-life, and incomplete knockdown. In addition, the organization of prokaryotic genes in operons and thus polarity effects cannot be overlooked as a biological consideration. As genome annotation of *S. epidermidis* improves, we anticipate that our group and others will be able to re-visit the data presented here for improved validation of this phenomenon and others. For the technical and biological reasons presented here, validation of high-quality hits through rigorous individual testing is still required before definitive conclusions can be drawn. Rather than serving as a conclusive study on gene function, we deem our CRISPRi data here to be a high throughput screening resource - as compared to individual knockout studies, the major advantage of this approach is its high-throughput format and the ability to quickly scale up to screen in multiple conditions, as performed.

Third, we note a major technical limitation to the CRISPRi approach in our non-model organism, which is the low transformability of this species. Future work should focus on increasing transformation efficiency or otherwise maneuvering around this roadblock. Indeed, there have been several advances in this field already that may be well applied for large-scale CRISPR/Cas targeting in non-model species (Brophy et al., 2018; Kristich et al., 2007).

In summary, our complementary approaches investigating gene fitness and gene plasticity across a diverse set of physiologically relevant conditions allowed us to make new biological insights into a keystone skin microbe. We also provided a real-world demonstration of both the promise and pitfalls of CRISPRi for high-throughput fitness profiling in non-model organisms. Future studies building on the accessibility and scalability of these approaches will further mechanistic investigation into the biodiversity of *S. epidermidis,* including the functional consequences of its extraordinary genetic diversity, as well as investigation into the tremendous biodiversity of its greater microbial neighbors.

## Methods

### CRISPRi Pooled Library Creation

#### Vector Design

Our CRISPR/dCas9 vector, which includes all components required for CRISPRi, has been previously validated(Spoto et al., 2020). The CRISPR/dCas9 vector is comprised of the dCas9 sequence (derived from pDB114dCas9 (Bikard et al., 2014) under the control of an anhydrotetracycline (ATc)-inducible promoter (derived from pRAB11(Helle et al., 2011)), a custom-designed dCas9 handle composed of a CRISPR RNA and a *trans*-activating small RNA fusion, two BSAI sites for ligation of sgRNA guides, and a chloramphenicol (Cm) resistance maker for selection.

#### sgRNA Design

sgRNAs 20 nucleotides in length were designed to target every NGG PAM site on both strands of the *S. epidermidis* Tü3298 genome using a customized version of the GuideFinder R scripts.

The Tü3298 strain of *S. epidermidis* was chosen for our screen for both technical and biological reasons. Firstly, it represents an interesting skin commensal strain due to its production of the lantibiotic epidermin (Augustin et al., 1992). Secondly, and importantly, the strain is amenable the phagemid plasmid transfer protocol (Winstel et al., 2016). Guides with GC content < 30% were discarded but no other filtering was performed after guide design. The identified sgRNA sequences were flanked with custom sequences containing BSAI restriction enzyme sites (for generation of single-stranded ends) and primer sites for PCR amplification according to the design protocol in SFig 4. In addition to 60,157 targeting guides, 100 scramble guides, were ordered from Twist Biosciences low-mass oligo pool. The 100 scramble guides were designed by blasting randomly generated sequences to the *S. epidermidis* Tü3298 genome. Only guides that did not match anywhere in the genome were selected as a scramble, non-targeting guide. All guides containing flanking sequences as ordered are available in the Supplemental Table 60.

#### Pooled library construction

The pool was amplified with custom PCR primers (F: 5’-GCTGTTTTGAATGGTCCC-3’ and R: 5’- CCGTTATCAACTTGAAAAAGTGG -3’) using NEB Q5 High-Fidelity DNA polymerase according to manufacturer’s instructions with an annealing temperature of 58C and an extension time of 30s for 10 cycles (Twist Biosciences recommends cycling between 6-12 times). The amplified pool was visualized on agarose gel, digested with BSAI to produce single-stranded overhangs and ligated into a BSAI digested pidCas9 targeting vector containing complementary single-stranded overhangs. To enhance transformation of our ligation library, ligation reactions (using NEB T4 ligase) were performed at 16C overnight in bulk and mixed in 0.5X volume with AMPure beads to remove impurities and concentrate the ligation 20X in nuclease-free water. The clean and concentrated ligation library was transformed into NEB 5alpha electro-competent *E. coli* and transformants were collected directly from the plate by flooding the plate with TSB/Cm. We estimate 250k colonies were recovered. The Qiagen Miniprep Kit was used for plasmid purification from the collected colonies, according to manufacturer’s instructions except that several lysate reactions were loaded on to one column sequentially prior to elution in nuclease free water. The isolated plasmids were concentrated to > 500 ng/uL prior to transformation into electrocompetent *S. aureus* PS187 to ensure a high number of transformants. *S. aureus* PS187 transformants were collected plate scraping and used directly for phagemid transfer. We estimate 115k colonies were recovered across approximately 100 plates. Plasmids were transferred to *S. epidermidis* Tü3298 from *S. aureus* PS187 via phagemid transduction as described elsewhere (Winstel et al., 2016). Briefly, bacteriophage Φ187 lysate and the plasmid-bearing *S. aureus* cells were incubated together for 30 minutes at 37 C without agitation and then moved to 30C with gentle agitation until the mixture cleared, indicating bacterial lysis by the phage, after approximately 3 hours. The phage lysate was centrifuged to pellet debris and filter sterilized twice with 0.22 micron filters prior to storage at 4C until use.

The phage lysate containing plasmid-bearing phage particles was incubated with *S. epidermidis* Tü3298 cells for 30 minutes and plated on TSA/chloramphenicol plates for selection of transformants. We estimate 150k colonies were recovered. Transformants were collected by the plate scraping method, diluted to an OD of ∼10 in TSB/Cm, and saved as glycerol stocks at -80C.

### CRISPRi Screening

#### Culture conditions

Our CRISPRi screening was conducted in 24 diverse stress and nutrient limiting conditions. The base media conditions are described as follows. For alternations to these base media conditions, including addition of stressors (e.g., hydrogen peroxide) see Table 1.

#### Defined media composition

Iscove’s Modified Dulbecco’s Medium (IMDM) with the following additions: 20 ug/mL Cm (for plasmid selection), 5g/L glucose, 20 mg/L adenine sulfate, 20 mg/L guanine HCL, 1:100 of MEM 100X Non-Essential Amino Acids Solution, 1:50 of MEM 50X Essential Amino Acids Solution, 25 uM FeCl3 (unless otherwise indicated, as in low and high iron conditions. pH adjusted to 7.0 with 1N NaOH.

#### Minimal Media composition

M9 Minimal Media according to Cold Spring Harbor protocol (Sander and Schneider, 1991), with the addition of FeCl3 to 25 uM, 20 ug/mL Cm (for plasmid selection), 2 mg/L thiamine HCL, and 2mg/L nicotinic acid. This solution was adjusted with varying concentrations of glucose (1% or 0.1% w/v) and amino acids (OmniPure CAA, 0.01% or 1% w/v)

#### CRISPRi growth assays

For each condition, media containing 0.1 uM aTC (for induction) and 20 ug/mL chloramphenicol (for plasmid selection) was inoculated 1:1000 with glycerol stock of the knockdown pool. The cultures were grown aerobically with a flask:media ratio of 5:1 at 37C with shaking, unless otherwise indicated. A list of tested conditions is available (Table 1). Two aliquots were taken at separate time points targeting early and early-mid exponential phase for each condition. Prior to each assay, 16-36 hour growth curves were determined for each condition using the Biotek Cytation Imaging Multi- Mode reader to estimate growth kinetics. After sampling, cells were spun down, the supernatant was removed, and the pellets were frozen at -20 C until preparation for sequencing

#### Small-scale Multiplexed CRISPRi

The 4-strain pool was made by mixing equal volumes of glycerol stock of previously created individual knockdown strains. For the 500 knockdown strain pool, guides were designed to gene body regions using a customized version of our GuideFinder script. Guides were ligated into our knockdown vector using the same approach for individual strain creation, as we previously described (Spoto et al., 2020). Briefly, guides were ordered as two single-stranded oligos (forward and reverse strands). Oligos were phosphorylated, annealed, and ligated into our BSAI-digested pidCas9 targeting vector. The procedure for transformation into *E. coli*, *S. aureus*, and phagemid transfer into *S. epidermidis* is the same as described for the large-scale pools. Culture conditions for both small knockdown pools are the same as described for the large pool analyses.

#### CRISPRi sequencing

For each condition, frozen cell pellets were washed in 1XPBS, subject to boiling alkaline lysis in 100mM NaOH at 95C for 5 minutes, and neutralized with 1M Tris 8.0 prior to PCR amplification off the crude lysate. Custom PCR primers were designed to work with the Illumina sequencing system to amplify the sgRNA region of the vector via primer binding sites on the pidCas9 vector up and downstream of the guide region (see STables 60-61 and SFig. 4). for primer design scheme and full list of primers used for sequencing). In brief, custom primers contain an adapter sequence, a Nextera index sequence for unique dual indexing (UDI), a sequencing primer, a stagger to maintain sequence diversity, and a PCR primer sequence (targeting up or downstream of the sgRNA guide sequence) for amplification of the product prior to sequencing. PCR was performed with NEB Q5 High-Fidelity DNA polymerase according to manufacturer’s instructions with an annealing temperature of 58C and an extension time of 30s for 25 cycles. The PCR product was visualized by agarose gel electrophoresis, the concentration was assessed with Qubit flourometric quantitation using a negative PCR reaction as a baseline and adjusted to 4nM with nuclease-free water. Samples were pooled and cleaned with AMPure beads (sequential 1.6x and .9x bead concentration cleaning steps) prior to sequencing. Sequencing was performed on the Illumina HiSeq and NovaSeq platforms targeting a read depth of ∼10-20M reads / sample.

#### Guide hit identification

Reads were trimmed to the sgRNA sequence using cutadapt (v1.18) (Martin, 2011) and aligned to a database containing all possible guides with BLASTN (NCBI blast+ v2.6.0) (Altschul et al., 1990) against a database of all designed sgRNA sequences (the read was considered a match with 100% identity and 100% coverage). Based on our design (described above), each guide was placed into one of the following categories based on their targeting location: 1) targeting a gene on the non-template strand 2) targeting a gene on the template strand 3) targeting a non-protein-coding sequence or 4) targeting multiple genomic locations. We expected that guides targeting genes on the non-template strand will lead to more profound gene knockdown (although differences in guide efficacy affect extent) as compared to guides targeting the template strand. We assumed guides targeting the template strand will not lead to fitness defects of vital genes. We assumed guides targeting the non-template strand of vital genes will often lead to fitness defects and acknowledge that not every guide will be effective in knocking down a gene. The raw read counts were input into a custom R script and we used DESeq2 (v1.26.0) (Love et al., 2014) to calculate a log2FC for each guide; default median of ratios normalization method was used. The adaptive shrinkage estimator was used for log2FC shrinkage via method=ashr (Zhu et al., 2019).

The log2FC reflects the difference in the relative abundance of the guide from each sampled time point to the initial pool. This analysis was performed to identify hits for each condition unless otherwise indicated (e.g., comparing induced versus uninduced samples for rich media pool characterization). To determine guides that were significantly reduced in the sampled time points as compared to the initial pool (“guide hits”), an FDR cutoff for each specific condition was determined (FDR that corresponds to the top 25% smallest FDR values). To be considered a “guide hit” a guide must meet this FDR threshold and have a log2FC < -1. Based on literature and our own experience, which indicates that not every guide targeted to a vital gene on the non-template strand will result in a fitness defect, we did not require that every guide (or any percentage of total guides) targeting a particular gene on the non-template strand be identified as a “guide hit” in order for the gene to be considered vital for growth. Along the same lines, we opted not to use median log2FC of guides targeted to a gene to determine gene fitness.

#### Investigation of essential and non-essential genes in rich media

To investigate the utility of our CRISPRi knockdown pool, we screened our pool in plain rich media (TSB) and plotted the distribution of log2FC values for guides targeting essential and non-essential genes. Our set of validating essential and non-essential genes were determined by homology (>30% homology over >90% sequence length) to known essential and non-essential genes in *S. aureus* (Sander and Schneider, 1991). Each *S. aureus* gene with more than one hit in the *S. epidermidis* was interrogated independently and the best hit was selected manually. Although there are likely some differences in essentiality between *S. aureus* and *S. epidermidis*, we believed there to be enough consistency in essentiality between the species to indicate whether our screen was feasible in this preliminary investigation.

#### Essential gene hit identification

Essential gene hits were binned according to high, medium, and low confidence as follows. High confidence genes are those that are identified by at least two “guide hits” in greater than ⅔ of all conditions. Medium confidence genes are those that are either 1) identified by at least two “guide hits” in ½ of all conditions or 2) identified by one “guide hit” in 2/3 of all conditions. Low confidence genes are those that are identified by at least one “guide hit” in ½ of all conditions. Gene accumulation curves for high, medium, and low confidence essential genes across conditions were generated using the vegan R package (v.2.5.7) (Okansen et al., 2020). One guide from each of these high confidence gene hits was randomly selected for individual knockdown strain creation and was created and individually assayed as described above in *CRISPRi growth assays*.

#### Conditional gene hit identification

Owing to the different growth rate of *S. epidermidis* across conditions and the level of growth that each of these conditions supports (a density plot of the log2FC values across the conditions (Fig. 2a) demonstrates this variation), we determined it was not appropriate to compare absolute log2FC values across conditions (i.e. we do not suggest that a gene has lower fitness in one condition vs another based on the degree of log2FC of the guides). To determine condition-specific gene hits, we identified hits for each condition (described above) and removed any putative essential genes that we identified with medium or high confidence. Thus, “condition-specific” gene hits are not those that are *exclusively* identified in one condition, but are those that are not generally identified across conditions (i.e., are not a putative essential gene).

#### Multi-stress response gene identification

To find multi-stress response genes, we searched for hits across multiple stress conditions in rich media. It is likely that nutrient availability affects gene fitness under stress and so we restricted our analysis to stress screens performed in rich media to reduce the effect of background media condition. Within our rich media stress conditions. These included the subinhibitory antibiotic (RM- mupirocin, RM-ciprofloxacin, RM-vancomycin), hyperosmotic (RM-sucrose, RM-glycerol, RM-urea, salt), pH (RM-acidic, RM-alkaline), heat (RM-42), sodium hypochlorite (RM-HOCl), and hydrogen peroxide (RM-H202) stress conditions. A gene was considered a “multi-stress response” gene if it was: 1) Identified as a hit (see above; briefly, by two or more independent guides with log2FC < -1) in 6 or more of these 13 stress conditions 2) Not identified as a hit in plain rich media (TSB) and 3) Not identified as a putative essential gene (medium or high confidence bin)

### Droplet-based CRISPRi

#### Droplet generation and growth assay

We piloted a droplet-based CRISPRi assay in plain rich media (TSB) and acidic rich media (RM-acidic; pH = 4.8). RainDrop Droplet Digital PCR system by RainDance Technologies was used to generate droplets containing the growth media with single *S. epidermidis* cells bearing the CRISPR/dCas9 vector. We calculated the number of *S. epidermidis* cells necessary to generate primarily empty droplets to avoid capture of multiple cells (∼ 2x10^4 cells /uL) and we confirmed the presence of single-cell droplets by light microscopy (SFig 3a-c). After droplet capture, the cultures were incubated at 37C and aliquots were taken periodically to assess growth of the cells in the droplets. Based on our batch culture results, we anticipated that growth in RM-acidic would be much slower than growth in plain RM. RM samples were grown for 8h; acidic-RM samples were grown for 20h; cultures were observed throughout via sampling and light microscopy to determine appropriate timing. At the end of each assay, droplets were broken by the addition of 1H,1H,2H,2H-Perfluoro-1-octanol. Amplicons were generated and sequenced as described above (“*CRISPRi sequencing*”) with the following modification: after droplet lysis, the oil phase was removed and spun down to pellet cells. Pellets were frozen at -80 until use.

#### Acid stress specific gene hits identification

Acid stress specific “guide hits” were determined as described above for our batch CRISPRi screen. Genes in this condition were identified as a hit if they met all three of the following conditions: 1) the gene was identified as a hit by two or more guides 2) the gene was not identified as a putative essential gene (by medium or high confidence) and 3) the gene was not identified as a hit by 2 or more guides in the RM droplet condition. We included this last constraint in our droplet-based CRISPRi screen to control for potential differences in gene fitness in droplet vs batch rich media (Okansen et al.).

## Transcriptomic Analysis

### Culture conditions

For each condition, media containing 0.1 uM aTC was inoculated 1:1000 with glycerol stock of the empty (non-targeting) vector. When possible, the same batch of media was used for the genome-wide screen assay and the RNA-seq assay and the samples were grown concurrently. The cultures were grown aerobically with a flask:media ratio of 5:1 at 37C with shaking, unless otherwise indicated. Two aliquots were taken at separate time points for each condition targeting early and mid-exponential phase. When necessary, cultures were moved into smaller flasks after sampling to maintain the 5:1 flask:media ratio. Cells were spun down, washed in 1XPBS, resuspended in Trizol and froze at -80C until RNA extraction

### RNA extraction, library preparation, and sequencing

RNA samples were extracted using the Qiagen RNeasy Plus Mini kit according to manufacturer’s instructions with the following modifications: Samples were thawed on ice, centrifuged to pellet and to remove Trizol supernatant, and resuspended in RLT Plus buffer with .1mm glass beads (autoclaved 2x prior to use). Samples were bead-beat using the Qiagen TissueLyser for 3 minutes and centrifuged to pellet beads. The supernatant was mixed with an equal volume of 70% ethanol. This mixture was applied to the Qiagen RNeasy column, and washed with appropriate buffers according to manufacturer’s instructions. An on-column DNAse digestion was performed according to the Qiagen RNeasy protocol using Qiagen RNAse-free DNAse. Samples were eluted in nuclease-free water according to manufacturer’s instructions. RNA was frozen at -80C until sequencing preparation. All RNA isolation steps were performed in a sterile hood following protocol to reduce RNA degradation (i.e. use of disposable plasticware/glassware, 2x autoclaved glass beads, RNaseZap RNAse decontamination solution use on all surfaces). RNA quality was analyzed using Agilent TapeStation and quantified with Qubit according to manufacturer’s instructions. RIN values ranged from 6.0-8.5 with a mean and median of 7.7. RNA was prepared for sequencing using the NEBNext rRNA Depletion Kit (Bacteria) and NEBNext Ultra II Directional RNA Library Prep Kit for Illumina kit according to kit instructions and sequenced on the Illumina NovaSeq, targeting 40-60M reads/sample. Sequences from each sample were trimmed of adapters using Trimmomatic (v0.32) (Bolger et al., 2014) and rRNA, tRNA, phiX sequences, and host contaminants were filtered out using BWA-MEM (v0.7.7)(Li and Durbin, 2009). BWA-MEM (v0.7.7). was used to align clean reads to reference Tü3298 genome. Mapped reads were assigned using featureCounts(v1.6.0) to generate a count table of reads to coding sequences (Liao et al., 2014).

### Differentially expressed gene identification

DESeq2 was used to identify differentially expressed genes, based on our input count tables. Unless otherwise indicated, plain rich media (TSB) was set as the reference for any stress conditions conducted in rich media. The plain defined media (DM) condition was set as the reference for any stress screen in this condition. M9 minimal media supplemented with 1% glucose and 1% Casamino acids was set at the reference for any screen conducted in M9 minimal media (concentrations of glucose and/or casamino acid additions vary). Log2FC values were calculated with the standard median of ratios normalization method; the adaptive shrinkage estimator was used for log2FC shrinkage via method=ashr (Zhu et al., 2019). Principle Coordinate Analysis (PCA) plotting of normalized read counts was used as a QC check to determine the relationship of the samples to one another. During this QC process, we found that the “mid-exponential” time points (all 3 biological replicates) for the plain rich media (TSB) condition clustered away from the remaining samples. Based on our analysis, we believe that these rich media samples are closer to the stationary phase and determined that it would not be appropriate to compare mid-exponential phase time points for the remaining samples. As such, we determined that it would be more appropriate to focus our remaining analysis on just the early-exponential time point for all of our samples. Using the shrunken log2FC values obtained from DESeq2, differentially enriched and repressed KEGG pathways were inferred using the GAGE package (v2.36.0). Pathview (v1.26.0) was used to generate visualizations of significantly enriched and repressed pathways as determined by GAGE.

## Supporting information

Stable1

Stable53

STables56&57

STable59

STable60

STable61

STables2to4

STables5to28

STables30to52

STables54to55

## Data Availability

All raw count data for and log2FC with fdr-adjusted p-values for every guide by CRISPRi screening is available in the Supplemental Data. Condition-specific hits and putative essential genes identified by CRISPRi screening are available as Supplemental Tables. Raw count data and log2FC with fdr-adjusted p-values for RNA-sequencing is available in the Supplemental Data and Tables respectively. pidCas9 vector design, Tu3298 annotation, PCR and sequencing primers are available in the Supplemental Data. Custom R script for guide design is available in the Supplemental Data and custom scripts for analysis can be made available upon request. All raw and processed sequencing data generated in this study will be available in the Short Read Archive (currently in submission SUB9556342)

## Author Contributions

We thank members of The Jackson Laboratory for Genomic Medicine (JAX-GM) Microbial Genomics core service, particularly Mark Adams, Joseph Brown, Sai Lek, and Purva Vats, for their assistance with the RNA-sequencing element of this project and the JAX-GM Genomic Technology core for their assistance with all sequencing components. We thank the members of the Oh laboratory for critical feedback on this project. We thank David Bikard and Luciano Marraffini for pDB114dCas9 and Ralph Bertram for pRAB11.

This work was funded by the National Institutes for Health (DP2 GM126893-01 and K22 AI119231-01). M.S. is funded by the National Institutes of Health (1F30DE027870-01 and T90-DE022526). J.O. is additionally supported by the NIH (1U54NS105539, 1 U19 AI142733, 1 R21 AR075174, and 1 R43 AR073562), the Department of Defense (W81XWH1810229), the NSF (1853071), the American Cancer Society, and Leo Foundation.

J.O defined the need for large-scale investigations into *S. epidermidis* gene function and provided guidance for the duration of the project. M.S. wrote the guide design script, built and screened the CRISPRi library, and designed CRISPRi/RNA-sequencing experiments. M.S., J.R, D.K., E.F., Y.O., and C.G. conducted bench worked related to pidCas9 vector design, multiplexed CRISPRi pilot studies, individual gene knockdown analyses, and RNA extraction.

M.S. analyzed the data. M.S. and J.O. drafted the manuscript. All authors read and approved the final manuscript. We declare that we have no competing interests.

---

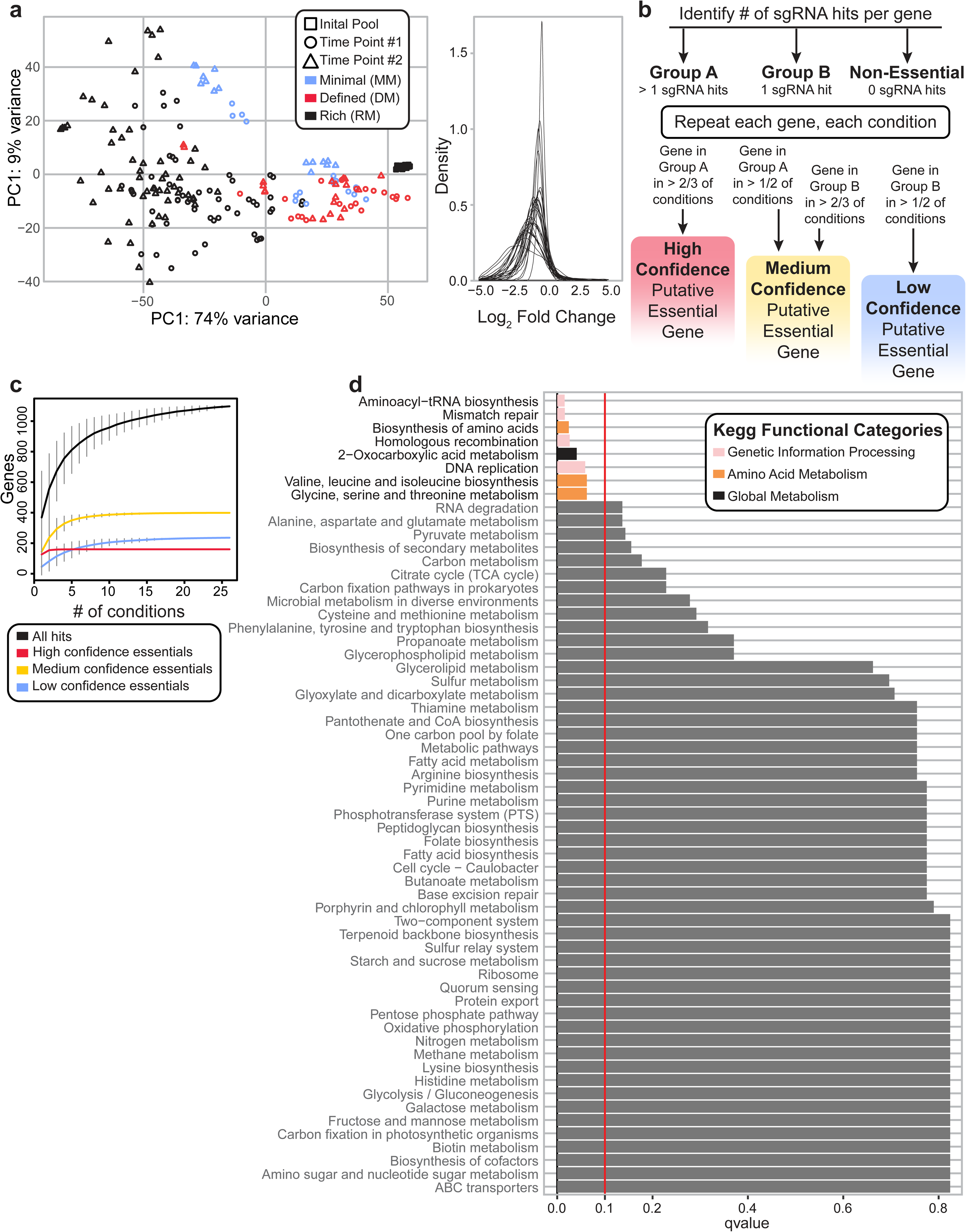

## References

A report from the NNIS System (2004). National Nosocomial Infections Surveillance (NNIS) System Report, data summary from January 1992 through June 2004, issued October 2004. American Journal of Infection Control 32, 470–485.

Altschul, S.F., Gish, W., Miller, W., Myers, E.W., and Lipman, D.J. (1990). Basic local alignment search tool. J Mol Biol 215, 403–410.

Asad, N.R., de Almeida, C.E.B., Asad, L.M.B.O., Felzenszwalb, I., and Leitão, A.C. (1995). Fpg and UvrA proteins participate in the repair of DNA lesions induced by hydrogen peroxide in low iron level in Escherichia coli. Biochimie 77, 262–264.

Augustin, J., Rosenstein, R., Wieland, B., Schneider, U., Schnell, N., Engelke, G., Entian, K.D., and Götz, F. (1992). Genetic analysis of epidermin biosynthetic genes and epidermin-negative mutants of Staphylococcus epidermidis. Eur J Biochem 204, 1149–1154.

Batzilla, C.F., Rachid, S., Engelmann, S., Hecker, M., Hacker, J., and Ziebuhr, W. (2006). Impact of the accessory gene regulatory system (Agr) on extracellular proteins, codY expression and amino acid metabolism in Staphylococcus epidermidis. Proteomics 6, 3602–3613.

Bikard, D., Jiang, W., Samai, P., Hochschild, A., Zhang, F., and Marraffini, L.A. (2013). Programmable repression and activation of bacterial gene expression using an engineered CRISPR-Cas system. Nucleic Acids Res 41, 7429–7437.

Bikard, D., Euler, C., Jiang, W., Nussenzweig, P.M., Goldberg, G.W., Duportet, X., Fischetti, V.A., and Marraffini, L.A. (2014). Development of sequence-specific antimicrobials based on programmable CRISPR-Cas nucleases. Nat Biotechnol 32, 1146–1150.

Bojanovič, K., D’Arrigo, I., and Long, K.S. (2017). Global Transcriptional Responses to Osmotic, Oxidative, and Imipenem Stress Conditions in Pseudomonas putida. Appl. Environ. Microbiol. 83.

Bolger, A.M., Lohse, M., and Usadel, B. (2014). Trimmomatic: a flexible trimmer for Illumina sequence data. Bioinformatics 30, 2114–2120.

Bosi, E., Monk, J.M., Aziz, R.K., Fondi, M., Nizet, V., and Palsson, B.Ø. (2016). Comparative genome-scale modelling of Staphylococcus aureus strains identifies strain-specific metabolic capabilities linked to pathogenicity. PNAS 113, E3801–E3809.

Brophy, J.A.N., Triassi, A.J., Adams, B.L., Renberg, R.L., Stratis-Cullum, D.N., Grossman, A.D., and Voigt, C.A. (2018). Engineered integrative and conjugative elements for efficient and inducible DNA transfer to undomesticated bacteria. Nature Microbiology 3, 1043–1053.

Byrd, A.L., Deming, C., Cassidy, S.K.B., Harrison, O.J., Ng, W.-I., Conlan, S., Belkaid, Y., Segre, J.A., and Kong, H.H. (2017). Staphylococcus aureus and S. epidermidis strain diversity underlying human atopic dermatitis. Sci Transl Med 9.

Canonaco, F., Schlattner, U., Wallimann, T., and Sauer, U. (2003). Functional expression of arginine kinase improves recovery from pH stress of Escherichia coli. Biotechnology Letters 25, 1013–1017.

Carey, A.F., Rock, J.M., Krieger, I.V., Chase, M.R., Fernandez-Suarez, M., Gagneux, S., Sacchettini, J.C., Ioerger, T.R., and Fortune, S.M. (2018). TnSeq of Mycobacterium tuberculosis clinical isolates reveals strain-specific antibiotic liabilities. PLoS Pathog 14, e1006939.

Casiano-Colón, A., and Marquis, R.E. (1988). Role of the arginine deiminase system in protecting oral bacteria and an enzymatic basis for acid tolerance. Applied and Environmental Microbiology 54, 1318–1324.

Chang, W., Small, D.A., Toghrol, F., and Bentley, W.E. (2006). Global Transcriptome Analysis of Staphylococcus aureus Response to Hydrogen Peroxide. Journal of Bacteriology 188, 1648–1659.

Chatterjee, P., Jakimo, N., Lee, J., Amrani, N., Rodríguez, T., Koseki, S.R.T., Tysinger, E., Qing, R., Hao, S., Sontheimer, E.J., et al. (2020). An engineered ScCas9 with broad PAM range and high specificity and activity. Nature Biotechnology 38, 1154–1158.

Chattopadhyay, M.K., and Tabor, H. (2013). Polyamines Are Critical for the Induction of the Glutamate Decarboxylase-dependent Acid Resistance System in Escherichia coli. J Biol Chem 288, 33559–33570.

Chaudhuri, R.R., Allen, A.G., Owen, P.J., Shalom, G., Stone, K., Harrison, M., Burgis, T.A., Lockyer, M., Garcia-Lara, J., Foster, S.J., et al. (2009). Comprehensive identification of essential Staphylococcus aureus genes using Transposon-Mediated Differential Hybridisation (TMDH). BMC Genomics 10, 291.

Cheung, G.Y.C., Joo, H.-S., Chatterjee, S.S., and Otto, M. (2014). Phenol-soluble modulins – critical determinants of staphylococcal virulence. FEMS Microbiol Rev 38, 698–719.

Christiansen, M.T., Kaas, R.S., Chaudhuri, R.R., Holmes, M.A., Hasman, H., and Aarestrup, F.M. (2014). Genome-Wide High-Throughput Screening to Investigate Essential Genes Involved in Methicillin-Resistant Staphylococcus aureus Sequence Type 398 Survival. PLOS ONE 9, e89018.

Coe, K.A., Lee, W., Stone, M.C., Komazin-Meredith, G., Meredith, T.C., Grad, Y.H., and Walker, S. (2019). Multi-strain Tn-Seq reveals common daptomycin resistance determinants in Staphylococcus aureus. PLoS Pathog 15, e1007862.

Cogen, A.L., Yamasaki, K., Sanchez, K.M., Dorschner, R.A., Lai, Y., MacLeod, D.T., Torpey, J.W., Otto, M., Nizet, V., Kim, J.E., et al. (2010a). Selective antimicrobial action is provided by phenol-soluble modulins derived from Staphylococcus epidermidis, a normal resident of the skin. J Invest Dermatol 130, 192–200.

Cogen, A.L., Yamasaki, K., Muto, J., Sanchez, K.M., Alexander, L.C., Tanios, J., Lai, Y., Kim, J.E., Nizet, V., and Gallo, R.L. (2010b). Staphylococcus epidermidis Antimicrobial δ-Toxin (Phenol-Soluble Modulin-γ) Cooperates with Host Antimicrobial Peptides to Kill Group A Streptococcus. PLOS ONE 5, e8557.

Corvaglia, A.R., François, P., Hernandez, D., Perron, K., Linder, P., and Schrenzel, J. (2010). A type III-like restriction endonuclease functions as a major barrier to horizontal gene transfer in clinical Staphylococcus aureus strains. PNAS 107, 11954–11958.

Cui, L., Vigouroux, A., Rousset, F., Varet, H., Khanna, V., and Bikard, D. (2018). A CRISPRi screen in E. coli reveals sequence-specific toxicity of dCas9. Nature Communications 9.

Dhandayuthapani, S., Blaylock, M.W., Bebear, C.M., Rasmussen, W.G., and Baseman, J.B. (2001). Peptide Methionine Sulfoxide Reductase (MsrA) Is a Virulence Determinant in Mycoplasma genitalium. J Bacteriol 183, 5645–5650.

Donati, S., Kuntz, M., Pahl, V., Farke, N., Beuter, D., Glatter, T., Gomes-Filho, J.V., Randau, L., Wang, C.-Y., and Link, H. (2021). Multi-omics Analysis of CRISPRi-Knockdowns Identifies Mechanisms that Buffer Decreases of Enzymes in E. coli Metabolism. Cell Systems 12, 56–67.e6.

Fedtke, I., Kamps, A., Krismer, B., and Götz, F. (2002). The Nitrate Reductase and Nitrite Reductase Operons and the narT Gene of Staphylococcus carnosus Are Positively Controlled by the Novel Two-Component System NreBC. Journal of Bacteriology 184, 6624–6634.

Finn, S., Rogers, L., Händler, K., McClure, P., Amézquita, A., Hinton, J.C.D., and Fanning, S. (2015). Exposure of Salmonella enterica Serovar Typhimurium to Three Humectants Used in the Food Industry Induces Different Osmoadaptation Systems. Appl. Environ. Microbiol. 81, 6800–6811.

Forsyth, R.A., Haselbeck, R.J., Ohlsen, K.L., Yamamoto, R.T., Xu, H., Trawick, J.D., Wall, D., Wang, L., Brown-Driver, V., Froelich, J.M., et al. (2002). A genome-wide strategy for the identification of essential genes in Staphylococcus aureus. Mol Microbiol 43, 1387–1400.

Gerdes, S.Y., Scholle, M.D., Campbell, J.W., Balázsi, G., Ravasz, E., Daugherty, M.D., Somera, A.L., Kyrpides, N.C., Anderson, I., Gelfand, M.S., et al. (2003). Experimental Determination and System Level Analysis of Essential Genes in Escherichia coli MG1655. J Bacteriol 185, 5673–5684.

Ghssein, G., Brutesco, C., Ouerdane, L., Fojcik, C., Izaute, A., Wang, S., Hajjar, C., Lobinski, R., Lemaire, D., Richaud, P., et al. (2016). Biosynthesis of a broad-spectrum nicotianamine-like metallophore in Staphylococcus aureus. Science 352, 1105–1109.

Gilbert, L.A., Horlbeck, M.A., Adamson, B., Villalta, J.E., Chen, Y., Whitehead, E.H., Guimaraes, C., Panning, B., Ploegh, H.L., Bassik, M.C., et al. (2014). Genome-Scale CRISPR-Mediated Control of Gene Repression and Activation. Cell 159, 647–661.

Goerlich, O., Quillardet, P., and Hofnung, M. (1989). Induction of the SOS response by hydrogen peroxide in various Escherichia coli mutants with altered protection against oxidative DNA damage. J Bacteriol 171, 6141–6147.

Gray, J.E., Richardson, D.K., McCormick, M.C., and Goldmann, D.A. (1995). Coagulase-negative staphylococcal bacteremia among very low birth weight infants: relation to admission illness severity, resource use, and outcome. Pediatrics 95, 225–230.

Grim, K.P., Francisco, B.S., Radin, J.N., Brazel, E.B., Kelliher, J.L., Solórzano, P.K.P., Kim, P.C., McDevitt, C.A., and Kehl-Fie, T.E. (2017). The Metallophore Staphylopine Enables Staphylococcus aureus To Compete with the Host for Zinc and Overcome Nutritional Immunity. MBio 8.

Guan, N., and Liu, L. (2020). Microbial response to acid stress: mechanisms and applications. Appl Microbiol Biotechnol 104, 51–65.

Helle, L., Kull, M., Mayer, S., Marincola, G., Zelder, M.-E., Goerke, C., Wolz, C., and Bertram, R. (2011). Vectors for improved Tet repressor-dependent gradual gene induction or silencing in Staphylococcus aureus. Microbiology (Reading, Engl.) 157, 3314–3323.

Hengge-Aronis, R. (1996). Back to log phase: σS as a global regulator in the osmotic control of gene expression in Escherichia coli. Molecular Microbiology 21, 887–893.

Jacobs, M.A., Alwood, A., Thaipisuttikul, I., Spencer, D., Haugen, E., Ernst, S., Will, O., Kaul, R., Raymond, C., Levy, R., et al. (2003). Comprehensive transposon mutant library of Pseudomonas aeruginosa. PNAS 100, 14339–14344.

Jensen, P.A., Zhu, Z., and Opijnen, T. van (2017). Antibiotics Disrupt Coordination between Transcriptional and Phenotypic Stress Responses in Pathogenic Bacteria. Cell Reports 20, 1705– 1716.

Jiang, W., Bikard, D., Cox, D., Zhang, F., and Marraffini, L.A. (2013). RNA-guided editing of bacterial genomes using CRISPR-Cas systems. Nature Biotechnology 31, 233–239.

Kappes, R.M., Kempf, B., and Bremer, E. (1996). Three transport systems for the osmoprotectant glycine betaine operate in Bacillus subtilis: characterization of OpuD. Journal of Bacteriology 178, 5071–5079.

Komatsuzawa, H., Ohta, K., Labischinski, H., Sugai, M., and Suginaka, H. (1999). Characterization of fmtA, a gene that modulates the expression of methicillin resistance in Staphylococcus aureus. Antimicrob Agents Chemother 43, 2121–2125.

Kristich, C.J., Chandler, J.R., and Dunny, G.M. (2007). Development of a host-genotype-independent counterselectable marker and a high-frequency conjugative delivery system and their use in genetic analysis of Enterococcus faecalis. Plasmid 57, 131–144.

Lai, Y., Di Nardo, A., Nakatsuji, T., Leichtle, A., Yang, Y., Cogen, A.L., Wu, Z.-R., Hooper, L.V., von Aulock, S., Radek, K.A., et al. (2009). Commensal bacteria regulate TLR3-dependent inflammation following skin injury. Nat Med 15, 1377–1382.

Lai, Y., Cogen, A.L., Radek, K.A., Park, H.J., MacLeod, D.T., Leichtle, A., Ryan, A.F., Di Nardo, A., and Gallo, R.L. (2010). Activation of TLR2 by a Small Molecule Produced by Staphylococcus epidermidis Increases Antimicrobial Defense against Bacterial Skin Infections. J Invest Dermatol 130, 2211–2221.

Lambers, H., Piessens, S., Bloem, A., Pronk, H., and Finkel, P. (2006). Natural skin surface pH is on average below 5, which is beneficial for its resident flora. International Journal of Cosmetic Science 28, 359–370.

Lee, H.H., Ostrov, N., Wong, B.G., Gold, M.A., Khalil, A.S., and Church, G.M. (2019). Functional genomics of the rapidly replicating bacterium Vibrio natriegens by CRISPRi. Nature Microbiology 4, 1105–1113.

Li, H., and Durbin, R. (2009). Fast and accurate short read alignment with Burrows–Wheeler transform. Bioinformatics 25, 1754–1760.

Liao, Y., Smyth, G.K., and Shi, W. (2014). featureCounts: an efficient general purpose program for assigning sequence reads to genomic features. Bioinformatics 30, 923–930.

Linehan, J.L., Harrison, O.J., Han, S.-J., Byrd, A.L., Vujkovic-Cvijin, I., Villarino, A.V., Sen, S.K., Shaik, J., Smelkinson, M., Tamoutounour, S., et al. (2018). Non-classical Immunity Controls Microbiota Impact on Skin Immunity and Tissue Repair. Cell 172, 784–796.e18.

Liu, X., Homma, A., Sayadi, J., Yang, S., Ohashi, J., and Takumi, T. (2016). Sequence features associated with the cleavage efficiency of CRISPR/Cas9 system. Scientific Reports 6, srep19675.

Love, M.I., Huber, W., and Anders, S. (2014). Moderated estimation of fold change and dispersion for RNA-seq data with DESeq2. Genome Biology 15, 550.

Ly, A., Henderson, J., Lu, A., Culham, D.E., and Wood, J.M. (2004). Osmoregulatory Systems of Escherichia coli: Identification of Betaine-Carnitine-Choline Transporter Family Member BetU and Distributions of betU and trkG among Pathogenic and Nonpathogenic Isolates. J Bacteriol 186, 296– 306.

Martin, M. (2011). Cutadapt removes adapter sequences from high-throughput sequencing reads. EMBnet.Journal 17, 10–12.

Mashruwala, A.A., and Boyd, J.M. (2017). The Staphylococcus aureus SrrAB Regulatory System Modulates Hydrogen Peroxide Resistance Factors, Which Imparts Protection to Aconitase during Aerobic Growth. PLOS ONE 12, e0170283.

Molina, J., Peñuela, I., Lepe, J.A., Gutiérrez-Pizarraya, A., Gómez, M.J., García-Cabrera, E., Cordero, E., Aznar, J., and Pachón, J. (2013). Mortality and hospital stay related to coagulase-negative Staphylococci bacteremia in non-critical patients. Journal of Infection 66, 155–162.

Moran, J.C., and Horsburgh, M.J. (2016). Whole-Genome Sequence of Staphylococcus epidermidis Tü3298. Genome Announc 4.

Moran, J.C., Alorabi, J.A., and Horsburgh, M.J. (2017). Comparative Transcriptomics Reveals Discrete Survival Responses of S. aureus and S. epidermidis to Sapienic Acid. Front Microbiol 8, 33.

Naik, S., Bouladoux, N., Wilhelm, C., Molloy, M.J., Salcedo, R., Kastenmuller, W., Deming, C., Quinones, M., Koo, L., Conlan, S., et al. (2012). Compartmentalized control of skin immunity by resident commensals. Science 337, 1115–1119.

Oh, J., Byrd, A.L., Deming, C., Conlan, S., NISC Comparative Sequencing Program, Kong, H.H., and Segre, J.A. (2014). Biogeography and individuality shape function in the human skin metagenome. Nature 514, 59–64.

Oh, J., Byrd, A.L., Park, M., Kong, H.H., and Segre, J.A. (2016). Temporal Stability of the Human Skin Microbiome. Cell 165, 854–866.

Okansen, J., Guillaume Blanchet, F., Friendly, M., Kindt, R., Legendre, P., McGlinn, D., Minchin, P.R., O’Hara, R.B., Simpson, G.L., Solymos, P., et al. (2020). vegan: Community Ecology Package.

van Opijnen, T., Bodi, K.L., and Camilli, A. (2009). Tn-seq: high-throughput parallel sequencing for fitness and genetic interaction studies in microorganisms. Nat Methods 6, 767–772.

Peters, J.M., Colavin, A., Shi, H., Czarny, T.L., Larson, M.H., Wong, S., Hawkins, J.S., Lu, C.H.S., Koo, B.-M., Marta, E., et al. (2016). A Comprehensive, CRISPR-based Functional Analysis of Essential Genes in Bacteria. Cell 165, 1493–1506.

Qi, L.S., Larson, M.H., Gilbert, L.A., Doudna, J.A., Weissman, J.S., Arkin, A.P., and Lim, W.A. (2013). Repurposing CRISPR as an RNA-Guided Platform for Sequence-Specific Control of Gene Expression. Cell 152, 1173–1183.

Queck, S.Y., Jameson-Lee, M., Villaruz, A.E., Bach, T.-H.L., Khan, B.A., Sturdevant, D.E., Ricklefs, S.M., Li, M., and Otto, M. (2008). RNAIII-independent target gene control by the agr quorum-sensing system: insight into the evolution of virulence regulation in Staphylococcus aureus. Mol. Cell 32, 150– 158.

Ren, X., Yang, Z., Xu, J., Sun, J., Mao, D., Hu, Y., Yang, S.-J., Qiao, H.-H., Wang, X., Hu, Q., et al. (2014). Enhanced specificity and efficiency of the CRISPR/Cas9 system with optimized sgRNA parameters in Drosophila. Cell Rep 9, 1151–1162.

Rousset, F., Cui, L., Siouve, E., Becavin, C., Depardieu, F., and Bikard, D. (2018a). Genome-wide CRISPR-dCas9 screens in E. coli identify essential genes and phage host factors. PLoS Genet 14.

Rousset, F., Cui, L., Siouve, E., Becavin, C., Depardieu, F., and Bikard, D. (2018b). Genome-wide CRISPR-dCas9 screens in E. coli identify essential genes and phage host factors. PLOS Genetics 14, e1007749.

Rousset, F., Cabezas-Caballero, J., Piastra-Facon, F., Fernández-Rodríguez, J., Clermont, O., Denamur, E., Rocha, E.P.C., and Bikard, D. (2021). The impact of genetic diversity on gene essentiality within the Escherichia coli species. Nature Microbiology 1–12.

Salama, N.R., Shepherd, B., and Falkow, S. (2004). Global transposon mutagenesis and essential gene analysis of Helicobacter pylori. J Bacteriol 186, 7926–7935.

Sander, C., and Schneider, R. (1991). Database of homology-derived protein structures and the structural meaning of sequence alignment. Proteins 9, 56–68.

Scharschmidt, T.C., Vasquez, K.S., Truong, H.-A., Gearty, S.V., Pauli, M.L., Nosbaum, A., Gratz, I.K., Otto, M., Moon, J.J., Liese, J., et al. (2015). A Wave of Regulatory T Cells into Neonatal Skin Mediates Tolerance to Commensal Microbes. Immunity 43, 1011–1021.

Schlag, S., Fuchs, S., Nerz, C., Gaupp, R., Engelmann, S., Liebeke, M., Lalk, M., Hecker, M., and Götz, F. (2008). Characterization of the oxygen-responsive NreABC regulon of Staphylococcus aureus. J Bacteriol 190, 7847–7858.

Segger, D., Aßmus, U., Brock, M., Erasmy, J., Finkel, P., Fitzner, A., Heuss, H., Kortemeier, U., Munke, S., Rheinländer, T., et al. (2008). Multicenter study on measurement of the natural pH of the skin surface. International Journal of Cosmetic Science 30, 75–75.

Shabayek, S., and Spellerberg, B. (2017). Acid Stress Response Mechanisms of Group B Streptococci. Front. Cell. Infect. Microbiol. 7.

Singh, V.K., Singh, K., and Baum, K. (2018). The Role of Methionine Sulfoxide Reductases in Oxidative Stress Tolerance and Virulence of Staphylococcus aureus and Other Bacteria. Antioxidants (Basel) 7.

Song, L., Zhang, Y., Chen, W., Gu, T., Zhang, S.-Y., and Ji, Q. (2018). Mechanistic insights into staphylopine-mediated metal acquisition. PNAS 115, 3942–3947.

Spoto, M., Guan, C., Fleming, E., and Oh, J. (2020). A Universal, Genomewide GuideFinder for CRISPR/Cas9 Targeting in Microbial Genomes. MSphere 5.

Thibault, D., Jensen, P.A., Wood, S., Qabar, C., Clark, S., Shainheit, M.G., Isberg, R.R., and van Opijnen, T. (2019). Droplet Tn-Seq combines microfluidics with Tn-Seq for identifying complex single-cell phenotypes. Nature Communications 10, 5729.

Tong, X., Campbell, J.W., Balázsi, G., Kay, K.A., Wanner, B.L., Gerdes, S.Y., and Oltvai, Z.N. (2004). Genome-scale identification of conditionally essential genes in E. coli by DNA microarrays. Biochemical and Biophysical Research Communications 322, 347–354.

Voloshin, O.N., Vanevski, F., Khil, P.P., and Camerini-Otero, R.D. (2003). Characterization of the DNA damage-inducible helicase DinG from Escherichia coli. J Biol Chem 278, 28284–28293.

Vylkova, S. (2017). Environmental pH modulation by pathogenic fungi as a strategy to conquer the host. PLOS Pathogens 13, e1006149.

Waldron, D.E., and Lindsay, J.A. (2006). Sau1: a Novel Lineage-Specific Type I Restriction-Modification System That Blocks Horizontal Gene Transfer into Staphylococcus aureus and between S. aureus Isolates of Different Lineages. J Bacteriol 188, 5578–5585.

Walton, R.T., Christie, K.A., Whittaker, M.N., and Kleinstiver, B.P. (2020). Unconstrained genome targeting with near-PAMless engineered CRISPR-Cas9 variants. Science 368, 290–296.

Wang, T., Wei, J.J., Sabatini, D.M., and Lander, E.S. (2014). Genetic screens in human cells using the CRISPR/Cas9 system. Science 343, 80–84.

Wang, T., Guan, C., Guo, J., Liu, B., Wu, Y., Xie, Z., Zhang, C., and Xing, X.-H. (2018). Pooled CRISPR interference screening enables genome-scale functional genomics study in bacteria with superior performance. Nat Commun 9, 1–15.

Widhelm, T.J., Yajjala, V.K., Endres, J.L., Fey, P.D., and Bayles, K.W. (2014). Methods to generate a sequence-defined transposon mutant library in Staphylococcus epidermidis strain 1457. Methods Mol Biol 1106, 135–142.

Wilde, A.D., Snyder, D.J., Putnam, N.E., Valentino, M.D., Hammer, N.D., Lonergan, Z.R., Hinger, S.A., Aysanoa, E.E., Blanchard, C., Dunman, P.M., et al. (2015). Bacterial Hypoxic Responses Revealed as Critical Determinants of the Host-Pathogen Outcome by TnSeq Analysis of Staphylococcus aureus Invasive Infection. PLoS Pathog 11, e1005341.

Williams, M.R., Cau, L., Wang, Y., Kaul, D., Sanford, J.A., Zaramela, L.S., Khalil, S., Butcher, A.M., Zengler, K., Horswill, A.R., et al. (2020). Interplay of Staphylococcal and Host Proteases Promotes Skin Barrier Disruption in Netherton Syndrome. Cell Rep 30, 2923–2933.e7.

Winstel, V., Kuhner, P., Rohde, H., and Peschel, A. (2016). Genetic engineering of untransformable coagulase-negative staphylococcal pathogens. Nat Protoc 11, 949–959.

Withman, B., Gunasekera, T.S., Beesetty, P., Agans, R., and Paliy, O. (2013). Transcriptional Responses of Uropathogenic Escherichia coli to Increased Environmental Osmolality Caused by Salt or Urea. Infection and Immunity 81, 80–89.

Xu, Y., Zhao, Z., Tong, W., Ding, Y., Liu, B., Shi, Y., Wang, J., Sun, S., Liu, M., Wang, Y., et al. (2020). An acid-tolerance response system protecting exponentially growing Escherichia coli. Nature Communications 11, 1496.

Zhou, C., and Fey, P.D. (2020). The acid response network of Staphylococcus aureus. Current Opinion in Microbiology 55, 67–73.

Zhou, W., Spoto, M., Hardy, R., Guan, C., Fleming, E., Larson, P.J., Brown, J.S., and Oh, J. (2020). Host-Specific Evolutionary and Transmission Dynamics Shape the Functional Diversification of Staphylococcus epidermidis in Human Skin. Cell 180, 454–470.e18.

Zhu, A., Ibrahim, J.G., and Love, M.I. (2019). Heavy-tailed prior distributions for sequence count data: removing the noise and preserving large differences. Bioinformatics 35, 2084–2092.

