## Supplemental_Note for "Large-scale CRISPRi and transcriptomics of *Staphylococcus epidermidis* identify genetic factors implicated in commensal-pathogen lifestyle versatility"

*Guide features and scramble guides*

Guide design is an important aspect of CRISPR-Cas studies, particularly in large-scale approaches where rules of thumb dictate automated selection of guides when manual curation is not feasible. Despite the importance of these design constraints, the literature in this area is sparse, particularly for CRISPRi in prokaryotes. We thus analyzed the relationship of certain guide features (e.g., GC content, distance from transcription start site) to gene knockdown to better understand general rules of guide design in *S. epidermidis*. While this approach is not a direct measure of gene knockdown, our investigation serves as a preliminary analysis for understanding guide design constraints in staphylococci*.*

Guide GC percentage has been previously reported to affect Cas9 targeting efficiency, although reports in multiple species are not fully concordant (Liu et al., 2016; Ren et al., 2014; Wang et al., 2014). For example, in human cells, guides with moderate GC content (40-60%) demonstrated more efficient cleavage than guides with more extreme GC percentages (Liu et al., 2016) - the rationale for the 35% GC minimum in our guide design. For guides targeting known essential genes (as determined by homology to *S. aureus* essential genes, the most appropriate proxy at this time), we did not observe any statistically significant difference between the log2FC of guides of varying GC content. As we limited our design to guides with ≥ 35% GC content, we cannot comment on the efficacy of guides with lower GC values, but our results here suggest that stringent GC content thresholds may not be a necessary design constraint for staphylococci (SFig.1a).

Distance of the guide target from the transcription start site (TSS) within a gene has been reported to affect guide efficiency in CRISPRi studies as well as numerous transposon-based gene inactivation studies (Jacobs et al., 2003; Qi et al., 2013). In an *E. coli* CRISPRi screen, guides targeting the first 5% of the gene had a significantly lower median log2FC compared to guides targeting any other region of the gene body (Qi et al., 2013). However, we did not observe any statistically significant difference between median log2FC for varying distances from the TSS for guides targeting essential genes. (SFig.1b)

Based on previous studies, we included ~100 non-targeting guides in our pool to control for the effects of expression of the CRISPR-Cas machinery on strain growth rate (Rousset et al., 2018a; Wang et al., 2014). We anticipated that these guides could be used to normalize fitness scores within and across conditions. Importantly, we identified that the non-targeting scramble guides in our screen do not behave neutrally as anticipated. We observed a surprising increase in the log2FC of these guides (substantially above most guides in the pool), suggesting that guides targeted to any region of the genome result in a mild growth defect compared to the non-targeting guides (SFig. 2a-b). The dissimilar behavior of these scramble guides as compared to the remainder of the pool makes them unsuitable to use as controls. Instead, we relied on the DeSeq2 default median of ratios method for normalization.

Altogether, our results can advise the design and execution of future high-throughput CRISPRi screens. Broadly, our results suggest that the behavior of CRISPRi screens are likely species and/or strain-specific. This is in view of the fact that our data are not fully concordant with previous CRISPRi benchmarking in *E. coli,* particularly as related to guide design and the use of scrambled non-targeting guides (Rousset et al., 2018a). Specifically, by demonstrating no difference

between the log2FC of guides with varying GC content targeting putative essential genes, we suggest that stringent GC content thresholds may not be necessary for guide design in Staphylococci and we advise further work in this area. If true, the removal of this design constraint would significantly improve the number of potential guides available for targeting and aid the design of large-scale CRISPRi studies. Additionally, improvements on CRISPR/Cas technology – such as the generation of PAM-less Cas proteins – will further expand the accessibility of these studies (Chatterjee et al., 2020; Walton et al., 2020). Importantly, we also discovered that non-targeting scrambled guides, which are commonly used as controls, may not behave neutrally as expected and thus make poor normalization controls. We advise that the use of non-targeting guides as normalization factors be done with great caution and only after preliminary studies indicate that their use is appropriate. In our study, we relied on the default DeSeq2 median if ratio methods for normalization but postulate that normalization based on a selection of validated, non-essential genes may be a more appropriate method moving forward. And finally, we note a major technical limitation to the CRISPRi approach in our non-model organism, which is the low transformability of this species. Future work should focus on increasing transformation efficiency or otherwise maneuvering around this roadblock.
